# What does not kill a tumour may make it stronger: *in silico* Insights into Chemotherapeutic Drug Resistance

**DOI:** 10.1101/230318

**Authors:** Sara Hamis, Perumal Nithiarasu, Gibin G Powathil

## Abstract

Tumour recurrence post chemotherapy is an established clinical problem and many cancer types are often observed to be increasingly drug resistant subsequent to chemotherapy treatments. Drug resistance in cancer is a multipart phenomenon which can be derived from several origins and in many cases it has been observed that cancer cells have the ability to possess, acquire and communicate drug resistant traits.

Here, an *in silico* framework is developed in order to study drug resistance and drug response in cancer cell populations exhibiting various drug resistant features. The framework is based on an on-lattice hybrid multiscale mathematical model and is equipped to simulate multiple mechanisms on different scales that contribute towards chemotherapeutic drug resistance in cancer. This study demonstrates how drug resistant tumour features may depend on the interplay amongst intracellular, extracelluar and intercellular factors. On a cellular level, drug resistant cell phenotypes are here derived from inheritance or mutations that are spontaneous, drug-induced or communicated via exosomes. Furthermore intratumoural heterogeneity and spatio-temporal drug dynamics heavily influences drug delivery and the development of drug resistant cancer cell subpopulations. Chemotherapy treatment strategies are here optimised for various *in silico* tumour scenarios and treatment objectives. We demonstrate that optimal chemotherapy treatment strategies drastically depend on which drug resistant mechanisms are activated, and that furthermore suboptimal chemotherapy administration may promote drug resistance.

## 1 Introduction

Chemotherapy is one of the major anticancer therapies, it is widely used both by itself and as part of multimodality treatment strategies. In most cases chemotherapy is effective, however the existence, or the development, of chemotherapeutic drug resistance in tumours continues to be a major problem in chemotherapeutic treatments, often leading to tumour recurrence post treatment [1, 18, 24, 35, 40, 42, 63]. Clinical and experimental observations suggest that cancers are often increasingly drug resistant subsequent to chemotherapy exposure [45, 60, 63, 76] and moreover cancer cells have the ability to posses, acquire and communicate drug resistant traits, enabling them to survive in the presence of chemotherapeutic drugs [35]. The existence of drug resistant phenotypes in cancer cell populations significantly impacts the efficacy and successfulness of chemotherapy [17, 69, 79].

The emergence of drug resistant cancer cells in tumours results in multiple subpopulations comprising drug sensitive (S) and drug resistant (DR) cells [17]. Furthermore cancer cell populations may evolve according to Darwinian principles [71] and cells that acquire drug resistance during chemotherapy have been observed to be increasingly metastatic [45], consequently DR subpopulations can reach significant proportions despite initially accounting only for a small fraction of some cancer cell population [35]. S and DR subpopulations that coexist synergistically compete for resources such as space and nutrients [39, 71], this competition influences the tumour environment and yields intratumoural heterogeneity. In clinical cases where tumour eradication is implausible, chronic control treatments can be proposed in which tumours are continuously managed and prohibited from reaching lethal proportions [25, 35, 39], long-term chemotherapy treatments are however linked to high frequency drug resistance [41, 66]. Since DR subpopulations are more fit than S subpopulations to survive in the presence of drugs, repeated or prolonged chemotherapy administration may amplify this fitness differentiation. Indeed Duan et al. [17] performed *in vitro* and *in vivo* experiments to conclude that in absence of drugs, sensitive cells are more fit than drug resistant cells and conversely, in presence of drugs DR subpopulations dominate as S subpopulations are reduced [17]. Thus drug resistant cells may thrive in micro-environments containing chemotherapeutic drugs, and a large DR subpopulation may result in disease recurrence post chemotherapy [67]. Ensuring that the DR subpopulation does not dominate the S subpopulation is of importance as such an outcome would render the tumour uncontrollable by chemotherapy [39]. This suggests that deliberately maintaining a subpopulation of drug sensitive cells may constitute a strategic countermeasure in tumour control schemes [48]. Duan et al. investigated the plausibility of this proposed strategy *in vivo* by comparing two cell populations exposed to chemotherapeutic drugs [17]. The first of these cell populations comprised drug resistant cells only, and the second population contained a combination of both drug resistant and sensitive cells. Their study confirmed that the second, combined, cell population was controllable by chemotherapy for a longer time period than the first, drug resistant, cell population [17].

Drug resistance is a multipart phenomenon which can be derived from several origins, in fact a cancer cell or tumour may express drug resistance in various ways [35, 48, 64]. Drug resistance may arise due to micro-environmental or intrinsic cell factors [19] and cells can acquire drug resistance by for example amplifying drug target molecules, activating DNA-repair, inducing drug transporters or altering their drug metabolism [64]. Phenotypical variations in cells, such as drug resistance, can be inherited or acquired and further, cells may be resistant to one specific drug or to multiple drugs, the latter phenomenon is known as multidrug resistance (MDR) [33, 35, 41, 71]. Early work performed by Luria and Delbrück on bacteria indicated that virus resistant mutations occur independently of the virus itself, thus indicating the existence of primary virus resistance [37]. These findings have since been adapted to oncology [18, 35], and primary drug resistance, that is drug resistance that occurs independently of the drug presence, is an accepted phenomenon arising from cell mutations. However, drug presence has been demonstrated to speed up the development of DR subpopulations [74] and cancer cells may acquire drug resistance by altering their genetic or epigenetic structure in order to evade drug effects [18]. Such alterations are induced by drug presence and may include dislodging drug receptors or overexpressing and modifying target molecules [18]. Heat shock proteins (Hsps) are molecular chaperones, continuously present in eukaryotic cells, yielding cytoprotective cell effects [29, 45]. Via their chaperoning actions they enable cells, both healthy and cancerous, to adapt to extracellular variations and maintain homeostasis whilst subjected to external stresses such as, maybe most importantly, hyperthermia but also hypoxia and anoxia, toxins and the presence of harmful chemical agents such as chemotherapeutic drugs [29, 45, 65, 76]. In healthy cells, the upregulation of Hsps can protect cells from for example high temperatures [45], however in cancerous cells Hsp upregulation may protect cells from drug effects [45, 76], thus enabling cells to survive under otherwise lethal conditions [29]. By extension Hsps have been linked to resistance to chemotherapeutic drugs [45] such as cisplatin, doxorubicin [76] and bortezomib [65].

Typical chemotherapy drugs target cells in active cell cycle phases, thus quiescent cells parry drug effects [40] and similarly slow-cycling cells are intrinsically more drug resistant than fast cycling cells [13, 67] as they are more likely to evade drug attacks. Slow-cycling cells have been linked to cancer stem cell-like (CSC-like) cells [40], they are important drivers for tumours due to their increased drug-survival rate and ability to serve as reserve stem cells [13, 67]. CSC-like cells have been depicted to display various traits including being slow-cycling, migratory and non-adhesive [1]. Rizzo et al. [60] demonstrated *in vivo* in mouse tails that a subpopulation of CSC-like cells indeed may benefit from drug presence when competing for resources with other cell populations. Thus slow-cycling cells have been identified to reinforce tumours, hence to eradicate cancer cell populations containing a subpopulation of slow-cycling cells it is crucial to target both slow-cycling and fast-cycling cells [13]. Intracellular communication is vital for multicellular organisms and cells may communicate with each other using chemical signalling, direct physical contact or, as discussed here, sending and receiving exosomes [66, 81]. Out of these listed information mediators, exosomes are of particular interest as they are detectable, cell type-specific and able to travel long distances [66, 81]. This implies that they could potentially constitute therapeutic targets or biomarkers and thus be used to impede or signal cancer [66, 81]. Exosomes constitute subcellular “molecule-parcels” that cells may utilise to dispose of non-essential materials [81], however perhaps more interestingly, they also facilitate long-distance intercellular communication by transporting information from sender cells to recipient cells [14, 30, 66]. These molecule-parcels contain biomolecules such as proteins, mRNA and DNA which may provide recipient cells with information that can be used to alter phenotypical attributes, in order to increase fitness [81]. Exosomes are a type of extracellular vesicle (EV) [11] and recent studies have identified EVs as key players in cancer development as they can influence tumour growth and metastasis by communicating oncogenic information [81]. EVs have also been assumed to be a part of the process that converts non-malicious cells into cancerous, and of optimising the balance between CSCs and non-CSCs [81]. Thus in response to chemotherapeutic drugs, cancer cells may not only develop individual drug resistance, but furthermore they may render other cells drug resistant by secreting exosomes to communicate and share drug resistant traits [11]. Exosomes may induce both destructive and protective cell responses, in fact the role of EVs depends on the regarded scenario [20]. In this study pathogenic exosomes only are modelled.

Mathematical models of tumour growth and treatment response may further cancer research by contributing insight into tumour dynamics, elucidating and validating clinically and experimentally recognised phenomena and guiding *in vitro* and *in vivo* experiments [25, 27, 35, 39, 69, 71]. Computational approaches to simulate biological systems are an important part of theoretical biology and may provide insights into biological phenomena [23]. *in silico* experiments have the advantage of cheaply being able to reproduce biological systems that span long time periods faster than real-time [71] and they can be used to find optimal treatment scheduling [32, 77]. Various such mathematical models of tumour growth, treatment response and drug resistance have previously been proposed [7, 35, 36, 47, 50, 56, 62, 68]. Roose et al. presented a comprehensive review of models of avascular tumour growth [62] and Lavi et al. compiled an extensive report discussing previous work on mathematical models of drug resistance in cancer [35]. name a few such models, Monro et al. [39] presented a continuum model in which tumour growth follows Gompertzian dynamics and drug resistant mutations occur proportionately to the tumour growth rate, in accordance with Luria Delbrück models. They concluded that increased drug administration may in fact reduce the survival length of a patient. Sun et al. [69] investigated therapy-induced drug resistance using a set of stochastic differential equations, they highlighted microenvironmental contributions to drug resistance. Powathil et al. [56] used the Compucell3D framework [70] to investigate two coexisting subpopulations, specifically one fast-cycling and one slow-cycling, in the presence of drugs to demonstrate intrinsic drug resistance of slow-cycling cells. There currently exists a number of hybrid discrete-continuum mathematical models that account for the multiscale nature of cancer [58], these models can be used to study tumour behaviour in response to multimodality treatment schemes [3, 4, 21, 28, 31, 49, 59]. Several modelling attempts have been made to address the multiscale aspects of cell growth by incorporating details such as vascular dynamics, oxygen transport, hypoxia, cell division and other intracellular features in order to study tumour dynamics and treatment response [38, 46, 51, 80]. Recently, Powathil et al. [53, 54] developed a hybrid multiscale cellular automaton, integrating cell cycle dynamics and oxygen distribution to study cell cycle-based chemotherapy delivery in combination with radiation therapy. As an important step towards personalised medicine, Caraguel et al. [9] managed to create virtual clones of *in vivo* tumours in mice using multiscale hybrid modelling. The tumour growth of various mouse tumours successfully agreed with the tumour growth of their respective virtual clones. Details of other multiscale cancer models are available in a review by Deisboeck et al. [15].

In the present *in silico* study, we propose a hybrid multiscale mathematical model that incorporates multiple types of drug resistance. *in silico* experiments are performed in order to study chemotherapeutic drug response in heterogeneous cancer cell populations hosting various types of drug resistant phenotypes pre, peri and post chemotherapy.

## 2 Model and *in silico* Framework

Following previous work by Powathil et al. [54], tumour growth and drug dynamics are modelled with a cellular automaton (CA). An overview of the relevant multiscale hybrid modelling techniques can be found in a later paper by Powathil et al. [55]. In this study, we expand on this model to incorporate multiple types of mechanisms that elicit drug resistance in cancer cells. Specifically, the CA used in this study can be categorised as a hybrid multiscale on-lattice model [58], incorporating a non-uniform micro-environment, extracellular dynamics, intracellular dynamics, intercellular dynamics and various categories of drug resistance regarded on a cellular resolution. The CA model uses mechanistic partial differential equations (PDEs), ordinary differential equations (ODEs) extracted from a regulatory molecular network, as well as stochasticity and phenomenological rules formulated by observations from biological experiments and clinical reports. An overview of the model schematics are illustrated in Figure 1 and details are provided throughout this section. The CA extends in two spatial dimensions, specifically a 100 by 100 square grid is utilised to simulate a physical tissue slab of (2 mm)^2^. This agrees with biological dimensions and each grid point is occupied by a cancer cell, a blood vessel cross section or extracellular matrix [54]. At the start of the *in silico* experiment one initial cancer cell is planted at the centre of the grid, over time this cell divides to give rise to a population of cancer cells and eventually chemotherapeutic drugs are applied to the system. Blood vessels are non-equidistantly scattered across the grid, they are assumed stationary and perpendicular to the two-dimensional tissue slab. Thus blood vessel cross sections live on the grid, where they act as source points for oxygen and drugs.

**Figure 1:**
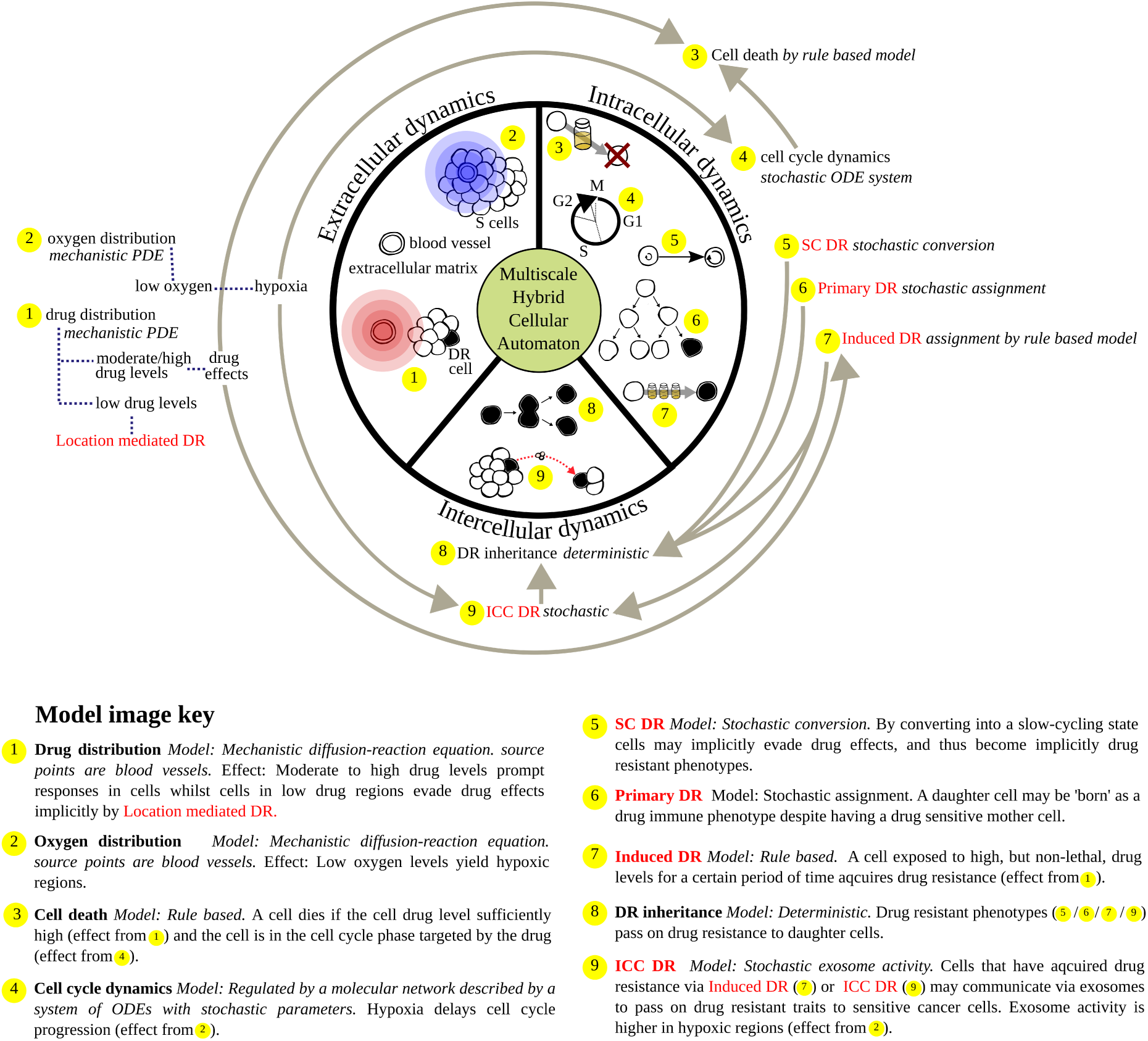
A schematic representation of the multiscale hybrid mathematical model used in this study. The model integrates extracellular, intracellular and intercellular dynamics. This is an on-lattice model and a lattice point may be occupied by a sensitive (S) or drug resistant (DR) cancer cell shown in black, a blood vessel cross-section or extracellular matrix. Various categories of drug resistance regarded on a cellular resolution are incorporated in the model, these categories are marked in red and listed in Table 1.

### 2.1 Intratumoural heterogeneity

Tumours are dynamic and should therefore be modelled as such [64], they are also heterogeneous and may constitute multiple distinguishable subpopulations [64, 71]. Intratumoural heterogeneity has been observed to promote drug resistance [64, 69] and hinder successful tumour prediction, thus by extension intratumoural heterogeneity complicates intelligent chemotherapy administration [71]. A tumour can express intratumoural heterogeneity in various ways and on multiple scales. For example nutrient concentrations, cell cycle dynamics and drug resistant traits may vary amongst cells in a tumour [69, 71]. Phenotypical attributes, such as drug resistant traits, may be acquired or inherited [71] and moreover stochasticity occurs naturally in biological processes. Hence various phenotypical subpopulations may arise in a cancer cell population, even if the population originates from one single cell [64]. To effectively treat tumours one should thus account for intratumoural heterogeneity, including the potential uprising of drug resistant subpopulations [71]. Our model accounts for intratumoural heterogeneity on various scales, details are provided in the following subsections. On a cellular level, each cell has an individual cell cycle length and individual drug resistant traits. On an extracellular level, the spatio-temporal micro-envrironment is highly dynamic, each cell has its own neighbours and moreover the blood vessels are non-equidistantly placed that oxygen and drug concentrations vary asymmetrically across the grid.

#### 2.1.1 Intracellular dynamics

The cell cycle mechanism can be partitioned into four sequential main phases, namely the gap 1 (G1), synthesis (Syn), gap 2 (G2) and mitosis (M) phase. In the synthesis phase, the DNA duplicates and in the mitoses phase chromosome segregation and cell division occur [26]. Between these two phases a cell requires time to grow and double its protein and organelle mass in order to replicate its DNA and divide, such intermediate phases are dubbed the gap phases. If some conditions are unfavourable a cell may delay its reproduction progress by exiting the cell cycle to enter an inactive, quiescent phase (G0) [54]. In the model, cancer cells are categorised as being in either the G1 phase or in the collective Syn-G2-M phase of the cell cycle, alternatively cells can exit the cell cycle and enter the quiescent phase G0 [54]. The cell cycle of each individual cell is governed by a regulatory molecular network described by a system of ODEs (Equation 1) [54] in which the dependent variables are protein concentrations and cell mass (*mass*). The relevant proteins are namely the Cdk-cyclin B complex (*CycB*), the APC-Cdh1 complex (*Cdh1*), the p55cdc-APC complex in its total form (*p*55*cdc_T_*) and active form (*p*55*cdc_A_*), and finally the Plk1 protein in its active form (*Plk1*). Details of protein functions and dynamics are provided in the Supplementary Material. The Cyclin B value is used as a marker to determine which state of the cell cycle that the cell is in, and by extension when cell division occurs as this happens when the cell exits Syn-G2-M phase to enter the G1 phase. Specifically, cell division occurs when the threshold value [*CycB*]_*thr*_ = 0.1 is crossed from above [54, 73]. When cell division occurs at time step *t_cd_* in the model, a daughter cell is placed on a grid point in the spherical neighbourhood of the parental cell, located in point *x_parent_*. At cell division the mass of the parent cell is halved so that 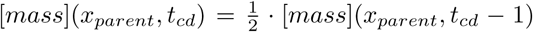 [73]. Grid points in lower order neighbourhoods are prioritised in this process, however up to third-level neighbourhoods are regarded. Each cell may divide until there is no unoccupied grid point on which to place a daughter cell, when this occurs a cell enters the quiescent phase G0. A cell may however re-enter the cell cycle if its neighbourhood is freed up and space is made available. Whilst in a quiescent phase cells are assumed to be drug immune in our model, this is because classical chemotherapy drugs target molecules that are overexpressed in specific cell cycle phases, for example the drug cisplatin affects the G1 phase of the cell cycle [54, 75]. The ODE that regulates the cell cycle reads

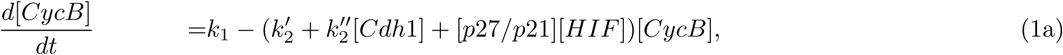

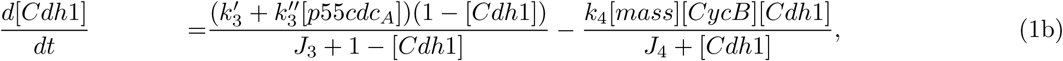

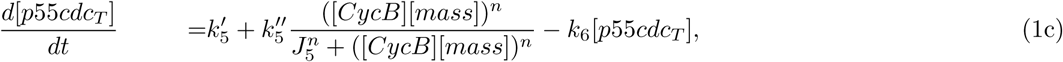

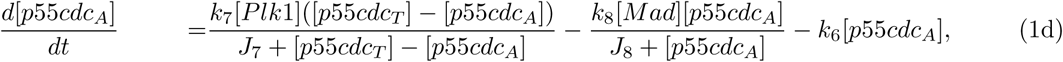

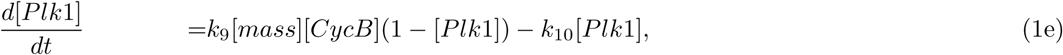

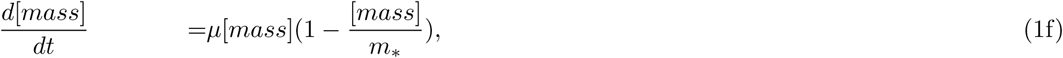

where the coefficient *μ* is individual to each cell and depicts cell growth rate, it takes a randomised value within an allowed range to account for heterogeneous cell cycle lengths amongst cells [54]. The variable [*HIF*] corresponds to the hypoxia inducible transcription factor-1 (HIF-1) pathway, in the model this factor is activated if the cell is classified as hypoxic [54] so that [*HIF*] = 1 in hypoxic cells and [*HIF*] = 0 in normoxic and hyperoxic cells. Oxygen distribution, and by extension hypoxia, is governed by an extracellular PDE, hence extracellular and intracellular dynamics are here integrated. Cellular accumulation of the HIF-1 *α* protein is furthermore associated with chemotherapeutic drug resistance [6]. Equation parameters are listed in the Supplementary Material.

#### 2.1.2 Extracellular dynamics

Extracellular dynamics is modelled using mechanistic PDEs describing oxygen and drug distribution. Oxygen is continuously produced at each time step on the blood vessels cross sections from which it is distributed across the grid according to the following diffusion-reaction equation [54],

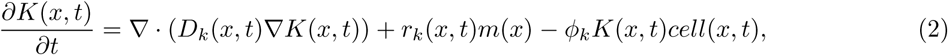

where *K*(*x, t*) denotes oxygen concentration in location *x* at time *t*, *D_k_*(*x, t*) is the oxygen diffusion coefficient, *r_k_* (*x, t*) is the oxygen production rate and *ϕ_k_* is the oxygen consumption rate. The variables *m*(*x*) and *cell*(*x, t*) are binary so that *m*(*x*) = 1 if there is a blood vessel in location *x* and *m*(*x*) = 0 otherwise. Likewise *cell*(*x, t*) = 1 if there is a cell in location *x* at time *t* and *cell*(*x, t*) = 0 otherwise. No-flux boundary conditions are applied, such boundary conditions coupled with the oxygen production at each time step will cause the total oxygen in the system to fluctuate over time. Thus in accordance with previous work by Powathil et al. [52], the absolute hypoxic threshold value will be different at each time step in the simulation, whilst the relative hypoxic threshold value will remain the same over time. This approach yields a spatial oxygen distribution at each time step which can be used to evaluate hypoxia. Physically, a cell is here classified as being hypoxic if its partial pressure of oxygen (pO_2_) is 10 mm Hg or less [10, 54]. Following Powathil et al. [54], a grid point in the implementation is defined to be hypoxic if it has a relative oxygen concentration of less than 0.1, where an oxygen concentration of 1 is normalised at the grid point with the highest oxygen concentration on the grid. Oxygen diffuses slower over grid points occupied by cancer cells than elsewhere and sufficiently high oxygen concentrations promote rapid cell proliferation, whilst hypoxia hinders cell cycle advancement. These hypoxic effects are incorporated in the model via the [HIF] parameter occurring in Equation 1 [54]. Chemotherapy drugs are similarly administered via blood vessel cross sections, however drugs are instantaneously produced at one single time step per drug administration. Drugs diffuse according to Equation 3,

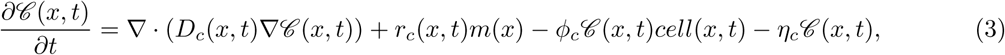

using no-flux boundary conditions. Here 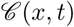 denotes drug concentration in location *x* at time *t*, *D_c_*(*x, t*) is the drug diffusion coefficient, *r_c_*(*x, t*) is the drug production rate, *ϕ_c_* is the drug consumption rate and *η_c_* is the drug decay rate. Chemotherapy drugs diffuse faster across extracellular matrix than inside the tumour [54]. Provided that the cell is in the drug-targeted cell cycle phase and that the cell is not explicitly drug resistant, a cell in location *x* at time *t* is killed if the drug concentration 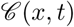 is such that 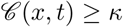, where *κ* is the lethal threshold drug concentration. When a cell dies, its grid point *x* becomes empty end the drug concentration 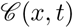 is halved, simulating drug binding by the cell. This drug binding compensates for the fact that the consumption term is omitted in Equation 3 as *ϕ_c_* is set to be 0, as is described in the Supplementary Material. Parameters occurring in the PDEs are also listed in the Supplementary Material.

### 2.2 Drug resistance

In this study, drug resistance is regarded on a cellular resolution, thus subcellular mechanics are simplified and drug resistance is categorised into cellular-level categories as illustrated in Figure 2. In the model drug resistance is firstly categorised as being either explicit or implicit. Explicit drug resistance occurs when a cell possesses any subcellular trait that directly protects it from drug effects, rendering it immune to some drug. Conversely, a cell displays implicit drug resistance when it is shielded from drug impact due to some indirect reason such as being slow-cycling or spatially located in a region of low drug concentration. Explicit drug resistant traits can be induced or inherited [34, 61] and also, cells may perform phenotypical alterations in response to intercellular interactions [64]. Therefore we here categorise drug resistance as being induced, primary or communicated. Induced drug resistance is activated in cells as a defensive response to drug presence whilst primary drug resistance is caused by cell mutations occurring independently of drugs [34, 61]. Cancer cells in which drug resistance has been induced may communicate and spread their drug resistant traits to other cancer cells via intracellular communication (ICC) in an effort to secure species survival [11, 66, 81]. Using hybrid modelling, the various categories of drug resistance are here modelled in different ways in order to easily reproduce biological phenomena in a way that is consistent with available clinical and experimental observations and data. In our model, drug resistance obeys rules formulated from previous findings from *in vitro* and *in vivo* results. Also incorporated are stochastic methods, as stochasticity occurs naturally in biological processes and may generate different cell phenotypes [64]. For all categories of phenotypical drug resistance, it is assumed that once a cell has established a drug resistant trait, its offspring will inherit that trait. This is in accordance with evolutionary Darwinian principles, as DR subpopulations are more fit to survive in drug presence than are S subpopulations [22]. Furthermore, the micro-environment influences drug transport across the tumour and impeded drug delivery by poor diffusion is indeed one of the primary reasons for treatment failure [35]. Thus drug efficacy and cytotoxic cell death is affected by the micro-environment, since drugs may not reach target cells. This may occur if the drug diffusion is impeded by dense population regions, or if the target cells are spatially located far away from drug source points, here blood vessels. To study how a heterogeneous micro-environment impacts drug resistance and drug response, blood vessels are non-equidistantly located across the *in silico* domain. Moreover, the speed of molecules such as oxygen, drugs and exosomes, depends on the medium that the molecules in question are traversing [54, 66]. This section provides information regarding the modelling of various drug resistant categories. Numerical values of the parameters introduced in this section are listed in the Supplementary Material, along with schematic representations of algorithms used in the model and a sensitivity analysis which demonstrates that our results are robust in regards to these parameters. Thus our qualitative findings, concerning drug response in cancer cell populations hosting various types of drug resistance, hold for variations of the chosen parameters.

**Figure 2:**
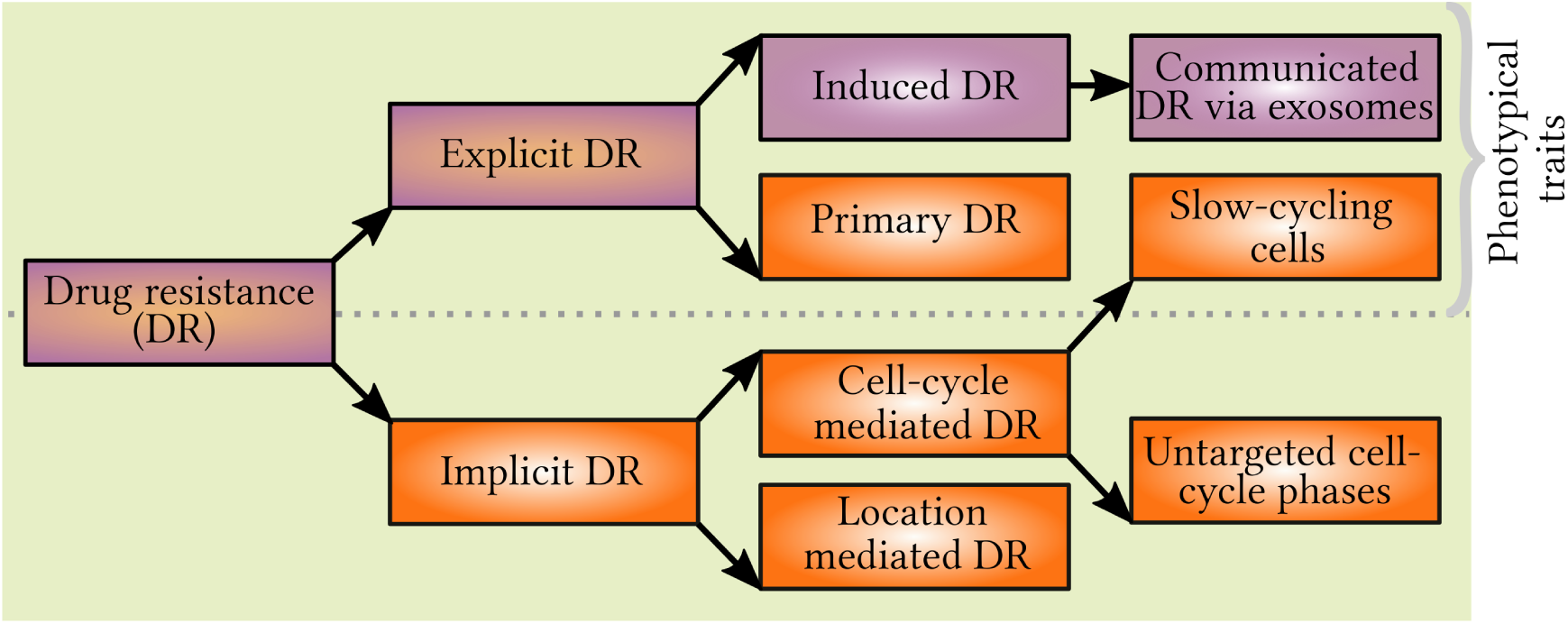
Classification of drug resistance categories occurring in the model. Drug resistance may be independent of drug presence (orange labels), induced as a consequence of drug presence (purple labels), or either (orange and purple labels). Cell acquisition of any DR phenotypical trait (top half) is here modelled as irreversible and inheritable by any future daughter cells to the cell in question.

#### 2.2.1 Primary drug resistance

Cells display primary drug resistance independently of drug presence, thus primary drug resistance may precede chemotherapy [35, 37, 61]. The *in silico* setup in our study is analogous to Luria and Delbrück’s [37] *in vitro* experiment, however here bacteria and virus have been exchanged for cancer cells and chemotherapy drugs respectively. Luria and Delbrück [37] stated that there was a probability per unit time that a sensitive cell would mutate into a, here drug, resistant phenotype. Thus in accordance with Luria and Delbrück models [37], primary explicit drug resistance is here modelled by stochastic cell mutations occurring at cell division. More specifically the chance of mutation is given by the mutation rate *α_pri_*, which corresponds to the probability of mutation per cell cycle. The parameter *α_pri_* is assumed to be small, so that drug resistant mutations are rare and moreover the probability that a mutated cell will revert back to a sensitive state is negligible and set to zero [37]. Hence in our model each daughter cell that is produced has a chance *α_pri_* of being explicitly drug resistance before being placed on the grid.

#### 2.2.2 Induced drug resistance

Drug presence has been demonstrated to speed up the development of DR subpopulations in cancers [74] and multiple studies have shown that cancer cells display altered epigenetic features following chemotherapy [34]. Cancer cells may become drug resistant after exposure to chemotherapy, the underlying cause of such induced drug resistance is assumed multifactorial [61]. Factors that may contribute to induced drug resistance include decreased apoptotic response, increased DNA-repair post drug-mediated damage, cell cycle alterations and reduced drug accumulation [61]. These factors may work concurrently and jointly towards establishing drug resistance in cells [61]. Hsps aid signalling pathways promoting cell growth and sustainability [45], and may induce anti-apoctic cancer cell properties [29]. Three members of the Hsp family that have been under scrutiny are namely Hsp27, Hsp70 and Hsp90, which have all been linked to promoting breast cancer tumours [45] and shown *in vivo* to contribute towards chemotherapeutic drug resistance [76]. In cancer cells, Hsps are plentiful and moreover administration of chemotherapeutic drugs has been observed to alter Hsp expression and increase Hsp activity [29]. Hsps may reside in both the cytoplasm and the cell nucleus, Vargas-Roig et al. demonstrated *in vivo* that chemotherapy drugs may modify the proportion of Hsps in different cell compartments [76], in fact drug administration resulted in increased nuclear expression, and decreased cytoplasmic expression of Hsp27 and Hsp70 [76]. Hsp27 promotes cell migration and differentiation *in vivo* [45] and elevated Hsp27 levels have been correlated to doxorubicon resistance and correspond to high tumorigenicity whilst low Hsp27 levels suppress tumour functions such as angiogenesis and proliferation of endothelial cells [45]. Hsp70 is linked to tumour growth and yields anti-apoptotic effect in tumours [45] and in breast cancer cells, a high propotion of nuclear Hsp70 is correlated to drug resistance [76]. Hsp90 is associated with regulating the cell cycle and controlling metastasis and proliferation [45] and high Hsp90 levels are linked to decreased survival rates in breast cancer patients [45]. Inhibiting Hsp27, Hsp70, Hsp90 has been hypothesised as part of future treatment plans [29, 65] and moreover Hsp70 has been suggested as a factor to prognostically evaluate the risk of disease recurrence [45]. The heat shock factor 1 protein (HSF1) regulates Hsps [65, 78], and HSF1 overexpression is linked to poor prognosis in breast, lung and colon cancers, an increase in intratumoural cancer stem cells proportions and chemotherapeutic drug resistance [78]. Specifically, cells with high HSF1 levels have displayed increased paclitaxel resistance [78]. Although cellular Hsp expression and activity have been linked to anti-cancer drug resistance, the details of these mechanisms are yet to be elucidated [76]. Thus in the mathematical model discussed in this paper, details concerning such underlying mechanisms of acquired drug resistance are omitted and we simply recognise the fact that if a cell has been exposed non-lethal drug levels for a certain amount of time, it can develop resistance to that drug. Furthermore, clinically, cancers are usually treated with combination therapies which makes it difficult to deduce rigorous information regarding how induced drug resistance is developed in cells as a response to one particular drug [72]. Due to this multifactorial nature of induced drug resistance, which involves various subcellular alterations occurring in response to drug presence in the micro-environment, in the model a cell obtains induced explicit drug resistance once it has been exposed to a high enough drug concentration for a sufficiently long time. Thus if a cell has experienced a minimum drug concentration *Xind* for *τ* time units, drug resistance is induced in the cell.

#### 2.2.3 Communicated drug resistance via exosomes

Srinivasan et al. [66] investigated exosome kinetics in lymphatic flow *in vitro* and *in vivo* using near infrared imaging. In the *in vitro* study, they found that planted exosomes from the HEY cell line, being spherical with a diameter of around 70 nm, travelled more effectively than size and density matched beads across lymphatic endothelial cells (LECs). This indicates that exosomes travel purposefully as opposed to randomly. Srinivasan et al. [66] reported an effective permeability for exosomes across the lymphatic endothelium in the order of 0.2 *μ*m/s and moreover exosomes were transported twice as fast across areas with cells compared to areas with no cells. Exosomes were observed to move rapidly *in vivo*, indeed they travelled 10 cm in a mouse tail within 2 minutes [66]. Studies also show that there is a correlation between the micro-environment and exosome activity, as exosome secretion and uptake is promoted in low-pH regions [81]. Further, hypoxic regions are associated with drug resistance [2] which is partly explained by elevated exosome secretion in such regions [11]. Here, exosomes are modelled using phenomenological rules formulated from experimental observations, incorporating stochasticity. Exosomes are modelled as discrete “molecule parcels”. Once per cell cycle each cell that has acquired drug or exosome induced drug resistance has a chance *α_ex_* of producing and secreting such a molecule parcel which is sent off in a random direction. The first sensitive cell that the exosome hits is dubbed the recipient cell, upon receiving the exosome the recipient consumes the parcel to gain drug resistance. Exosome production and uptake times are incorporated in the travel time and thus modelled as instant, using data from Srinivasan et al. [66] we choose a propagation speed of 0.2 *μ*m/s which is equivalent to 1 grid point (20 *μ*m) per 100 s when travelling across cells, and half of this speed when travelling across extracellular matrix. The chance of exosome production *α_ex_* is significantly higher in hypoxic regions in order to account for increased exosome activity in such regions [11].

#### 2.2.4 Cell cycle mediated drug resistance by slow-cycling cells

Slow-cycling cells are distinguishable *in vitro* [13] and Srinivasan et al. [67] demonstrated that fast and slow-cycling cells may coexist in a tumour. Many chemotherapeutic drugs, such as cisplatin, attack only cells in a certain phase of the cell cycle by targeting proteins overexpressed in the corresponding phase, leaving other cells unaffected [54, 75]. Since the half-life times of common chemotherapy drugs are significantly shorter than the average cell cycle length of standard eukaryotic cells [54], slow-cycling cells are implicitly more resistant to chemotherapy as they are likely to evade drug impact whilst being in a prolonged untargeted cell cycle phase. Consequently, if there exists a subpopulation of slow-cycling cells in a tumour, this subpopulation is more likely to survive chemotherapy and proliferate post treatment, despite having a slower production rate. Such a slow-cycling subpopulation may comprise CSC-like cells, as slow-cycling cells have been linked to cancer stem cells [40], which in turn have been conferred with reduced sensitivity to chemotherapeutic drugs [78]. Previous research indicates that cancer cells may obtain stem-cell like traits [16], in fact non-stem cancer cells may convert into CSC-like cells seemingly spontaneously, as demonstrated *in vivo* by Chaffer et al. [12]. Micro-environmental factors, such as oxygen supply, may influence such conversions however here cell conversion into a slow-cycling state is modelled as independent of the micro-environment. The [HIF] parameter, occurring in Equation 1, which is switched on in hypoxic cells only does however increase the cell cycle length of all cells, fast-cycling and slow-cycling, and thus in the model oxygen levels effect cell-cycle lengths. Slow-cycling cells are multidrug resistant as they are resistant to any drug that targets only a subset of cell cycle phases. In the model we assume that slow-cycling conversion occurs spontaneously, independently of drug presence. More specifically, once per cell cycle there is a chance *α_SC_* that a sensitive, fast-cycling cell will spontaneously convert into a slow-cycling state. Here slow-cycling cells are assumed to have a cell cycle length that is roughly twice as long as a sensitive cell [56]. The chance of conversion from a slow-cycling, implicitly drug resistant state back into a fast-cycling state is assumed negligible in accordance with previously discussed Luria Delbrück models [37] and Darwinian principles and thus set to zero. Furthermore the *in silico* experiment spans 700 hours only, and the chance that a cell converts and re-convert is assumed negligible in the model. Once a cell is randomly selected to convert into a slow-cycling state, the individual growth rate factor *μ* of the regarded cell, occurring in Equation 1, is updated in order to achieve slower cell-cycle progression and by extension a longer cell-cycle length. Daughter cells to slow-cycling cells are assigned appropriate growth rates associated with slow-cycling cells at creation. Permitted growth rates for both sensitive fast-cycling cells and slow-cycling cells are listed in the Supplementary Material.

### 2.3 Implementation

The CA is implemented in C/C++ using a high performance computational framework. Ordinary differential equations are solved using a fourth order Runge-Kutta method and partial differential equations are solved using explicit finite difference schemes. Prior to commencing the *in silico* experiment, the micro-environment is implemented and thus blood vessel cross sections are scattered in a non-equidistant fashion across the grid, this is in order to emphasise and study the effects of spatio-temporal oxygen and drug heterogeneity. One initial cancer cell is then planted at the centre of the grid, from this originating cell the cancer cell population will grow. Tumour growth is simulated over 700 hours, where it has been concluded that 700 hours is a sufficiently long simulation time to study the drug resistance and drug response in the system. Here, a time step size Δ*t* = 10^−4^ hours, is used in accordance with the appropriate nondimensionalisation of the oxygen parameters occurring in Equation 2 [54]. Cisplatin, a chemotherapy drug which here is modelled to attack G1 cells only, is administered in two instances at 500 and 600 hours. For each such instance, drugs are produced on all blood vessels at one single time step. The total amount of drug produced at one time step, on all blood vessels, corresponds to the drug dosage, which can be varied to study various cases. Specifically, here the studied drug dosage are: (1) 1C, (2) 2C, (3) 4C or (4) 8C, as listed in Table 2. The dosage 1C is parametrised so that, in the absence of drug resistant phenotypes amongst the cells, 1C kills half of the total cell population shortly after the first drug administration at 500 hours. By doubling the drug dosage once, twice or thrice to 2C, 4C or 8C respectively a higher cell kill is achieved. However, the relationship between the drug dosage and the number of cells killed is not linear, as is further discussed in Section 3.1.1. For the fixed chemotherapy schedule at 500 and 600 hours, the administered drug dosages are varied and the size of the sensitive and drug resistant cell subpopulations are computed pre, peri, and post chemotherapy. Cell-maps are produced in order to display cell population topology over time, these cell-maps are visualised in ParaView [5]. Five different *in silico* experiments are here performed, corresponding to five different categories of drug resistance namely (a) to (e) listed in Table 1. Each experiment is performed 100 times, subsequently average values and standard deviations are reported. A sensitivity analysis is performed in order to confirm that our results are robust in regards to small parameter variations. This sensitivity analysis is available in the Supplementary Material.

**Table 1:** The labelling of drug resistance categories occurring in the model.

| Label | Drug Resistance Category |
| --- | --- |
| a | No drug resistance (No DR) |
| b | Primary drug resistance (Primary DR) |
| c | Induced drug resistance (Induced DR) |
| d | Induced and, via exosomes, communicated drug resistance (ICC DR) |
| e | Cell cycle mediated drug resistance by spontaneous conversion to slow-cycling cells (SC DR) |

**Table 2:** The labelling of drug dosages used in the implementation. For non-zero drug dosages the labelling system is such that label *ℓ* corresponds to drug dosage 2*^ℓ−1^*.

| Label | Drug dosage |
| --- | --- |
| 0 | 0 C (No drug) |
| 1 | 1 C |
| 2 | 2 C |
| 3 | 4 C |
| 4 | 8 C |

## 3 Results and Discussion

### 3.1 Results

In absence of chemotherapeutic drugs, the growth of the DR subpopulation in our *in silico* experiment is proportional to the growth of the S subpopulation, as demonstrated by the graphs in Figure 3. This is in accordance with previous mathematical models and experimental results [35, 37, 39].

**Figure 3:**
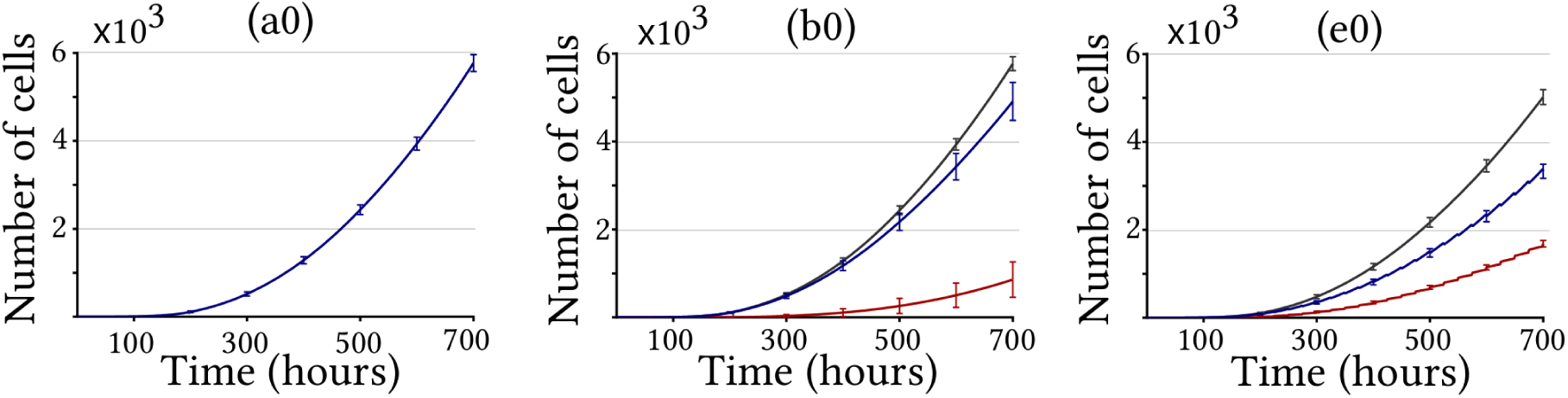
Growth of the cancer cell population over time in drug absence, showing total population (black graph), drug sensitive subpopulation (blue graph) and drug resistant subpopulation (red graph) incorporating drug resistance categories (a0) No DR, (b0) Primary DR and (e0) SC DR. Cases (c0) Induced DR and (d0) ICC DR are omitted since they produce the same results as (a0) in drug absence. Each graph shows the average number of cells based on 100 simulations, the standard deviation is demonstrated with error bars.

#### 3.1.1 No drug resistance

Once chemotherapy is applied to the *in silico* setup, the cancer cell population decreases in response to drugs. This is especially clear when no drug resistant phenotypes exist, the cancer cell population then rapidly reduces after drug administration. However population growth quickly resumes, and the size of such populations eventually reaches, and surpasses, the pre-chemotherapy population size. This is evident in graphs (a1) through to (a4) in Figure 4, and especially clear for high chemotherapy dosages where the size of the cell population cycles between being large immediately before drug administration and being small directly after drug administration. Rottenberg and Borst [63] demonstrated similar cyclic behaviour in mouse tails *in vivo*. Increasing the drug dosage trivially kills more cancer cells, however tumour topology and spatial heterogeneity significantly affects drug transport. Moreover, due to intratumoural heterogeneity in regards to the cell cycle, some cells will be shielded from the drug simply by being in an untargetted cell cycle phase. This multiscale heterogeneity affects the relationship between drug dosage and drug efficacy, where drug efficacy here corresponds to the number of cells killed by the drug. Graphs (a1) to (a4) in Figure 4 demonstrate the poor scaling between drug dosage and drug efficacy. Further, here we define the scaling efficiency *ϵ* by

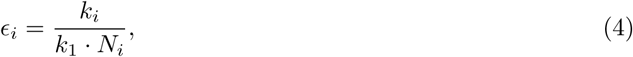

where *k_i_* denotes the number of cells killed in experiment *i* and *N_i_* denotes the dimensionless drug dosage coefficient used in experiment *i*. Thus here, *i* = 1, 2, 3, 4 and respectively *N_i_* = 1, 2, 4, 8. If the relationship between drug dosage and drug efficacy was linear, doubling the drug dosage would mean doubling the number of cancer cells killed. In such an ideal case it would hold that *ϵ_i_* = 1 ∀*i*, however this is not the case, as illustrated in Figure 5 which shows the scaling efficiency for the first drug administration at 500 hours. Hence in a clinical setting, the harm that may follow increased toxicity from higher drug dosages may not be worth the modest increase in killed cancer cells. The cell-maps in Figure 6 further highlight how the spatial heterogeneity affects drug delivery and consequently where the drug concentrations are high enough for cell kills to occur.

**Figure 4:**
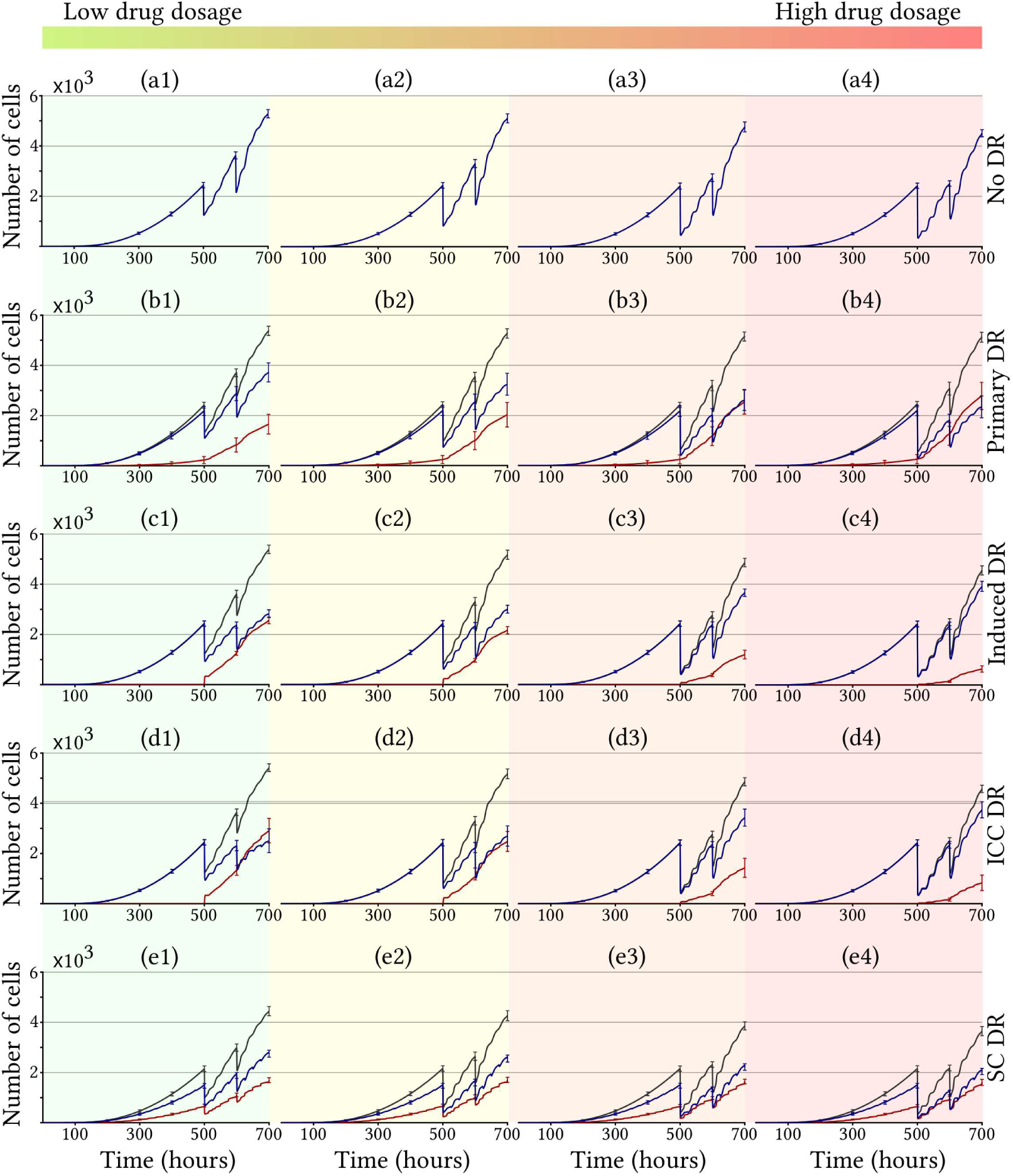
Growth of the cancer cell population over time when drugs are applied at 500 and 600 hours, showing total population (black graph), drug sensitive subpopulation (blue graph) and drug resistant subpopulation (red graph). Each row in the figure corresponds to a category of drug resistance (a) to (e) according to the labelling in Table 1, whilst each column corresponds do a specific drug dosage varying from low (leftmost column) to high (rightmost column), namely (1) 1C, (2) 2C, (3) 4C and (4) 8C according to the labelling system in Table 2. Each graph shows the average number of cells based on 100 simulations, the standard deviation is demonstrated with error bars.

**Figure 5:**
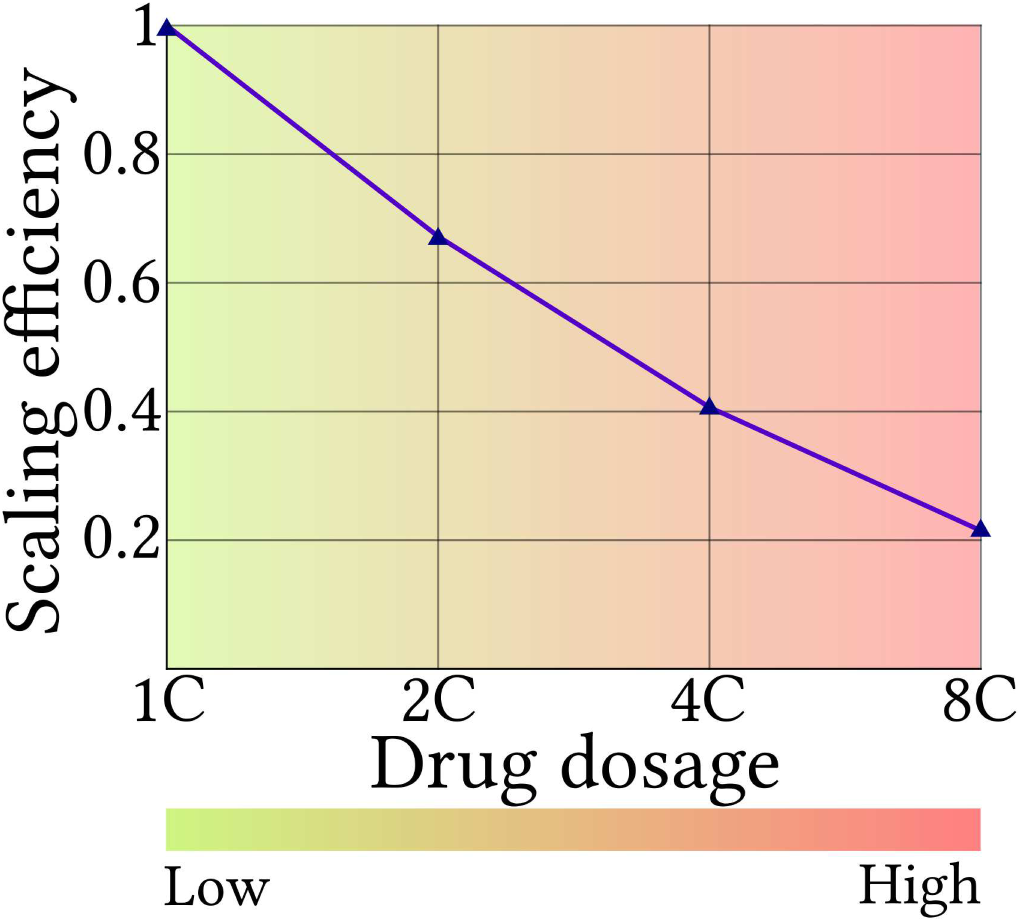
The scaling efficiency, demonstrating the relationship between drug dosage and drug efficacy in terms of number of killed cancer cells. This is for the first drug administration at 500 hours for experiments (a1) to (a4) according to the labelling in Tables 1 and 2, where no drug resistant phenotypes are present. Results are based on average values for 100 tests.

**Figure 6:**
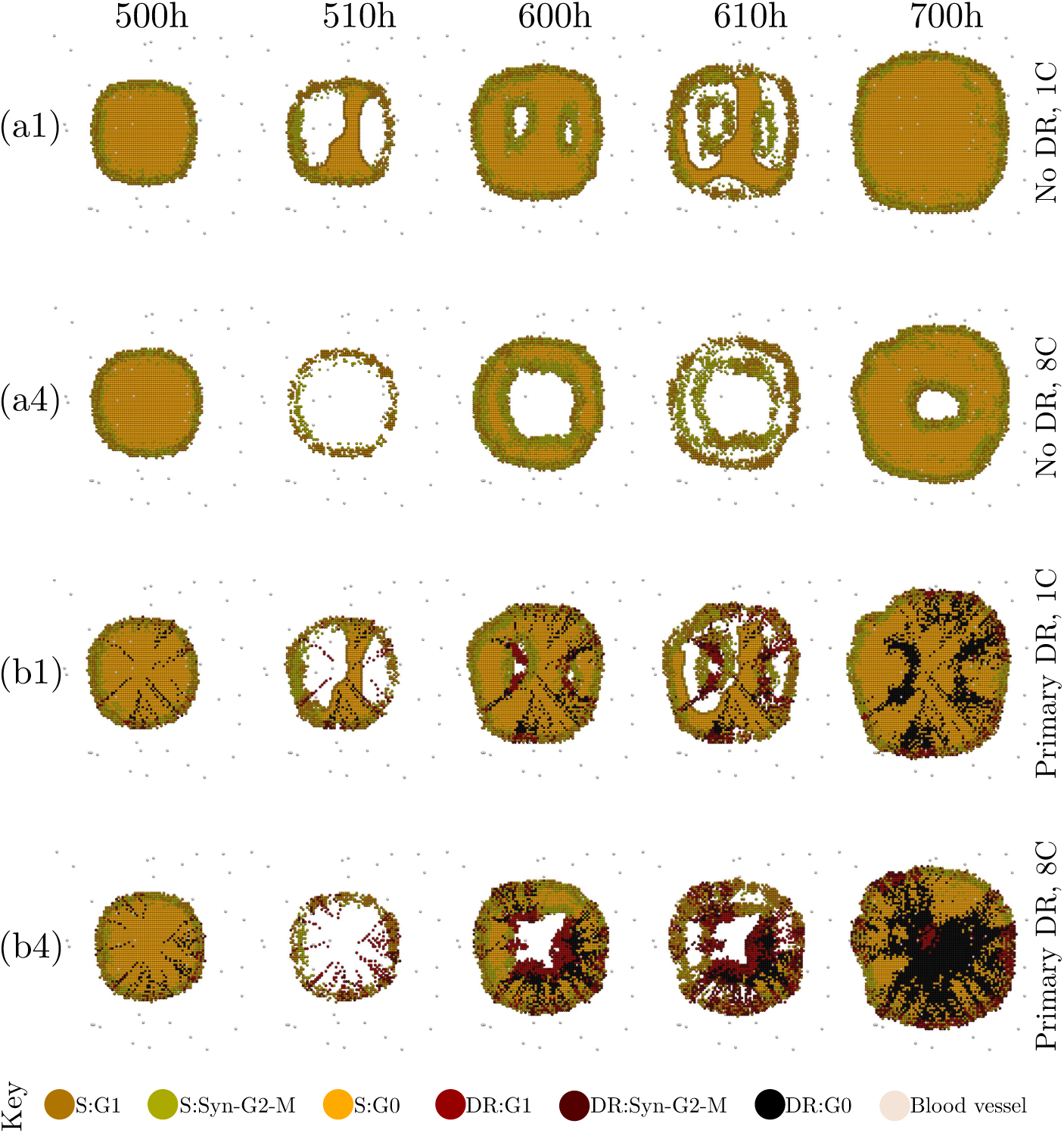
Cell-maps of the cancer cell populations at times 500 h (immediately before first drug dose), 510 h 600 h (immediately before second drug dose), 610 h and 700 h (end of simulation). Cases incorporating (a) No DR and (b) Primary DR are shown for drug dosages of (1) 1C and (4) 8C according to the labelling in Tables 1 and 2. White areas correspond to extracellular matrix.

#### 3.1.2 Primary drug resistance

When a cancer cell population expresses primary drug resistance, two subpopulations, comprising sensitive and drug resistant cells, already coexist prior to chemotherapy. Once administered, chemotherapy eliminates cells belonging to the S subpopulation whilst leaving the DR subpopulation unharmed. Hence increasing the drug dosage, and consequently killing more sensitive cells, enables the DR subpopulation to flourish as it gains access to more resources such as space and oxygen. Graphs (b1) through to (b4) in Figure 4 show that such DR subpopulations indeed benefit from high drug environments. Over all, our results show that chemotherapeutic treatments yield poor results if there exists a subpopulation of drug resistant cells prior to commencing treatment, this supports clinical observations depicting that primary drug resistance gravely reduces the successfulness of chemotherapy and influences the choice of treatment regime [18]. In our experiment, primary DR subpopulations grows outwards in radial strands pre chemotherapy, as illustrated in cell-maps (b1) and (b4) in Figure 6. This geometry is explained by the model, in which primary drug resistant mutations occur at cell division, prior to placing the cell on the grid. Moreover drug resistant cells produce drug resistant offspring, thus radial strands are formed as the cancer cell population grows. This DR subpopulation will spread from these strands, which consequently will widen post drug administration. On the other hand, subpopulation S will spread from regions containing sensitive cells that survived drug effect. Such sensitive survivor cells are clustered in regions of low drug concentrations, typically located far away from blood vessels and enclosed by other cancer cells, as drugs travel more slowly over dense population regions.

#### 3.1.3 Induced drug resistance

For simulations incorporating induced drug resistance, the DR subpopulation arises post chemotherapy. Here drugs diffuse more slowly over cancer cells than over extracellular matrix. Therefore, once the initial effect of chemotherapy has eliminated the majority of the sensitive cancer cells, and the tumour has disintegrated into clusters of surviving cancer cells, drug resistant cells are typically located on boundaries between cancer cells and extracellular matrix, as illustrated in cell-maps (c1) and (c4) in Figure 7. On these boundaries drug resistance is induced since the cells are exposed to high, but non-lethal, drug concentrations for a sufficiently long time. From these points of origin the DR subpopulation spread. By increasing the amount of chemotherapy, the cells that are exposed to this intermediate, high but non-lethal, drug concentration is reduced. Hence higher chemotherapy dosages do not only kill more cells overall, but reduces the amount of drug resistant phenotypes, as demonstrated by graphs (c1) to (c4) in Figure 4, this result applies to a cancer cell population in a confined space.

**Figure 7:**
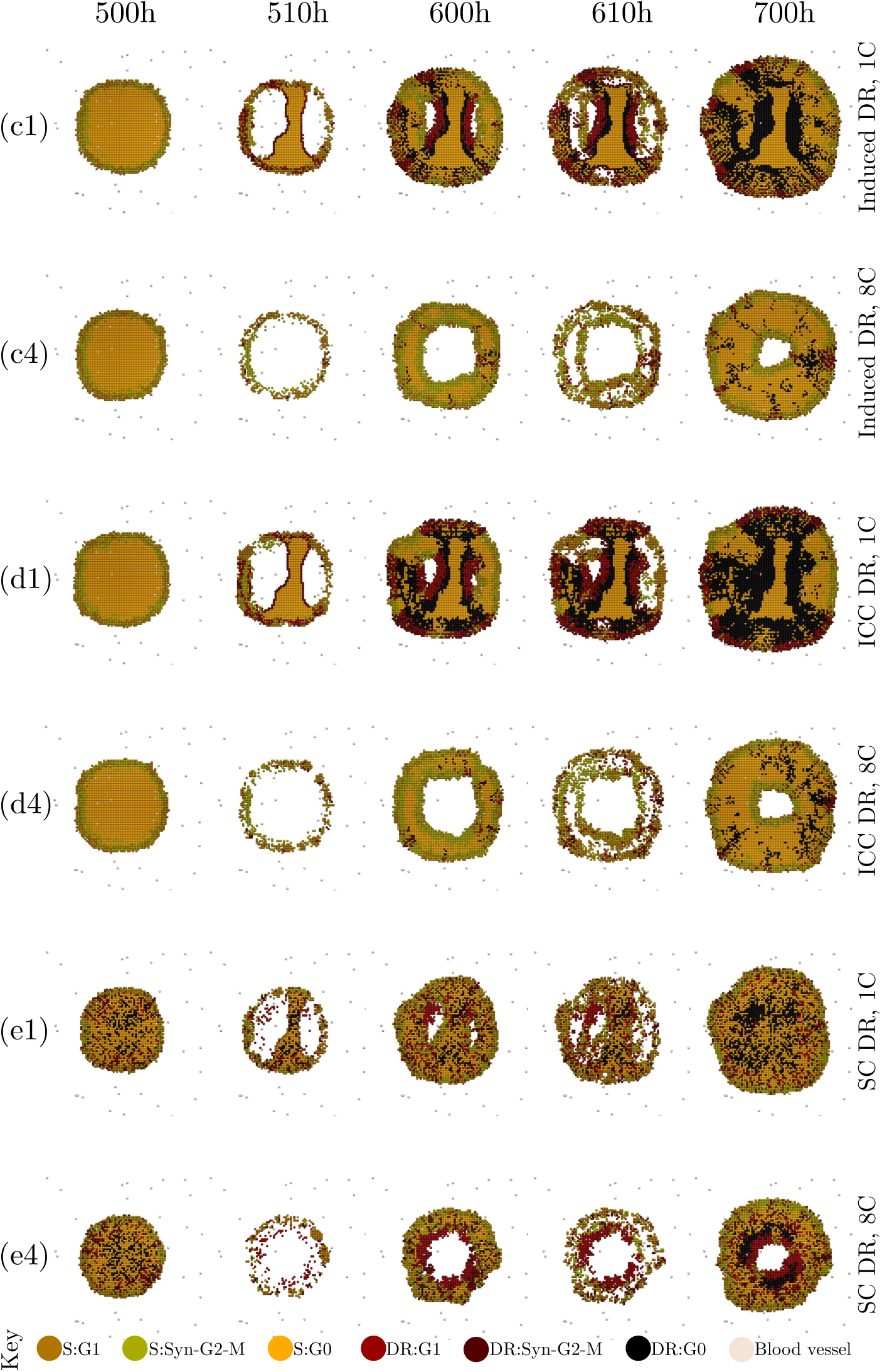
Cell-maps of the cancer cell populations at times 500 h (immediately before first drug dose), 510 h 600 h (immediately before second drug dose), 610 h and 700 h (end of simulation). Cases incorporating (c) Induced DR, (d) ICC DR and (e) SC DR are shown for drug dosages of (1) 1C and (4) 8C according to the labelling in Tables 1 and 2. White areas correspond to extracellular matrix.

#### 3.1.4 Communicated drug resistance via exosomes

The effect of ICC is demonstrated by comparing experiment (c) Induced drug resistance, to experiment (d) Induced and, via exosomes, communicated drug resistance. This can be done by regarding graphs (c1) to (c4) and (d1) to (d4) in Figure 4 and cell-maps (c1), (c4), (d1), (d4) in Figure 7. These figures show that the communicative exosome component amplifies the effect of induced drug resistance alone. The number of oncogenic exosomes produced increases with the number of drug resistant cells. In low to moderate drug environments, many exosomes are thus produced, as there exists many cells which have acquired induced DR in accordance with the results in Section 3.1.3. Conversely, in very high drug regimes fewer cells will survive to acquire induced drug resistance and consequently fewer oncogenic exosomes will be produced. Exosomes have been hypothesised as possible treatment targets, and our results indicate that reducing the exosome activity would aid the S subpopulation, as less cells would convert from sensitive to drug resistant. Here, hypoxic cells secrete more exosomes than do normoxic cells. This results highlights one of the benefits of targeting hypoxic tumour regions, as doing so may reduce exosome activity and by extension hinder communicated drug resistance.

#### 3.1.5 Cell cycle mediated drug resistance by slow-cycling cells

Slow-cycling cells are more likely to evade drug effects, as shown in graphs (e1) to (e4) in Figure 4 where, after each drug attack, the DR slow-cycling subpopulation displays a higher survivor rate. High chemotherapy dosages increase this effect, and thus benefit the DR subpopulation. Since the conversion to slow-cycling cells is modelled as spontaneous, implicit drug resistance may precede chemotherapy. Furthermore this spontaneity means that at every cell cycle, each fast-cycling cell has the same chance *α_SC_* of converting to a slow-cycling state. Thus pre chemotherapy the DR subpopulation will be point-wise scattered across the cell population, and post chemotherapy the DR subpopulation will spread from these source points, as illustrated in cell-maps (e1) and (e4) in Figure 7.

#### 3.1.6 Cell-map topology

Each drug resistance category in our model corresponds to a typical cell-map topology and vice-versa, as shown in Figures 6 and 7. The location of the subpopulations S and DR depends on the category of drug resistance regarded, as does the location of regions with high cell-kill numbers. These cell-maps are useful for conveying spatial heterogeneity, which is of importance since intratumoural heterogeneity is known to heavily influence drug efficiency [54]. Any sensitive cell, or cell cluster, that is surrounded by a band of other cells will be partly shielded from drug effects, as drugs travel slowly over dense population regions. Further, any quiescent sensitive cell, or cell cluster, that is surrounded by a band of explicitly drug resistant cells is eternally safe from drug presence in our *in silico* setup. This is because such a sensitive cell, or cell cluster, will never meet the necessary condition to re-enter the cell cycle as this requires free neighbourhood space. Thus even sensitive cell may be protected from drug effect, this is an example of location mediated implicit drug resistance.

### 3.2 Discussion

Our study shows that drug response in cancer cell populations is crucially influenced by the drug resistant phenotypes amongst the cells. *in silico* we demonstrate that the effect of chemotherapy is heavily dependent not only on the mere existence of drug resistant cells, but also the type of drug resistance displayed and micro-environmental factors. Clinically this implies that optimal chemotherapy scheduling and dosages depend on tumour specific data, including information regarding drug resistance and tumour environment. Indeed our results show that some types of drug resistant phenotypes thrive in low drug settings, whilst other flourish in high drug settings.

Before proceeding to discuss optimal drug dosages, one must define what constitutes as “successful” chemotherapy. Is the aim perhaps to (*i*) reduce the cancer cell population as much and quickly as possible or to (*ii*) be able to control the tumour long-term using chemotherapy? Case (*i*) may be relevant when chemotherapy is used in combinations with other treatments, for example when neoadjuvant chemotherapy is used prior to radiation treatment or surgery. Conversely, Case (*ii*) may be applicable when chronic chemotherapeutic treatment strategies are used, as can be done when it is implausible to completely eliminate a tumour. In such cases it is vital to suppress any DR subpopulation in order to keep the tumour manageable by chemotherapy. For Case (*i*), Figure 8 provides the data needed to discuss intelligent treatment strategy. This diagram trivially shows that high chemotherapy dosages are the most effective to rapidly eradicate cancer cells. However, what is not shown in our result, but is of clinical importance, is that high drug dosages are coupled with high toxicity which may be harmful to patients. Moreover our results show that the relationship between drug dosage and drug efficacy scales poorly, which is worth considering in a clinical setting. The increase in toxicity following from an increased drug dosage may not validate the outcome in terms of tumour reduction. In Case (*ii*) there is a balance, and sometimes a trade-off, between eliminating sensitive cells and aiding the drug resistant cells. Indeed killing of sensitive cells paves the way for any drug resistant phenotypes. Hence administering low chemotherapy dosages sometimes constitutes a wiser treatment strategy as it delays the uprising of a drug resistant subpopulation. Evolutionary theory and the interaction between S and DR subpopulations may thus play a role in designing intelligent treatment strategies. Figure 9 shows the ratio between S and DR subpopulations at different times. This diagram demonstrates that drug resistant phenotypes that arise independently of drug presence benefit from high drug dosages. However, *in silico*, the opposite is true here for drug-induced drug resistant phenotypes, which prosper in low drug conditions. Ideal chemotherapeutic tumour treatments would involve rapidly reducing tumour size whilst minimising drug resistance, thus meeting the requirements of both Cases (*i*) and (*ii*). However, our results indicate that when using chemotherapy only, there is a trade-off between tumour reduction and the suppressing of drug resistant phenotypes in some cases, thus the objectives of Cases (*i*) and (*ii*) may conflict.

**Figure 8:**
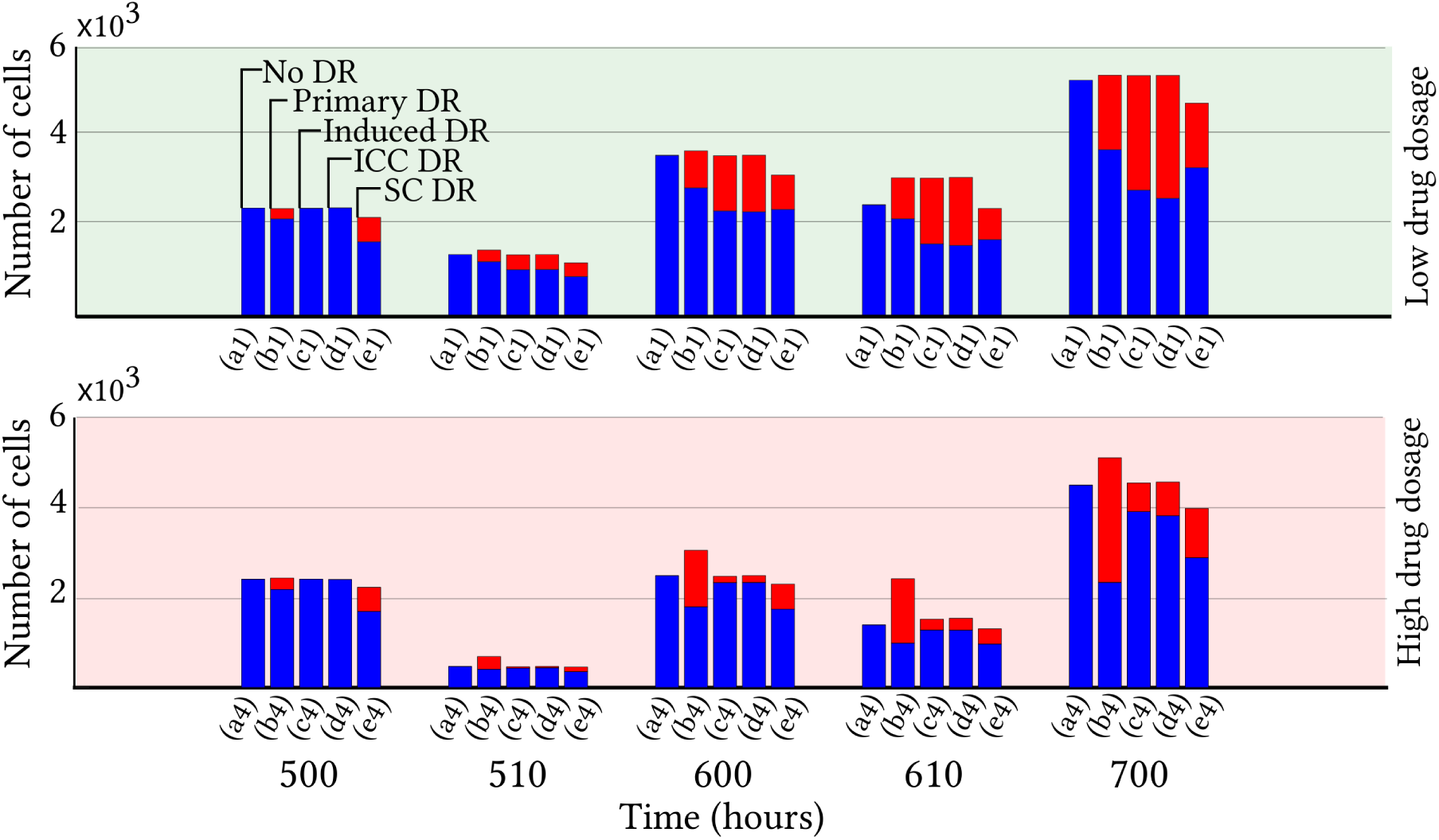
Diagram of the drug sensitive subpopulation (blue) and the drug resistant subpopulation. Various types of drug resistance are incorporated namely (a) No DR, (b) Primary DR, (c) Induced DR, (d) ICC DR, (e) SC DR. Results are shown at times 500 h (immediately before the first drug dose), 510 h, 600 h (immediately before the second drug dose), 610 h and 700 h (end of simulation) for low drug dosages, (1) 1C, and high drug dosages, (4) 8C according to the labelling in Table 2. Each diagram shows the average value based on 100 simulations.

**Figure 9:**
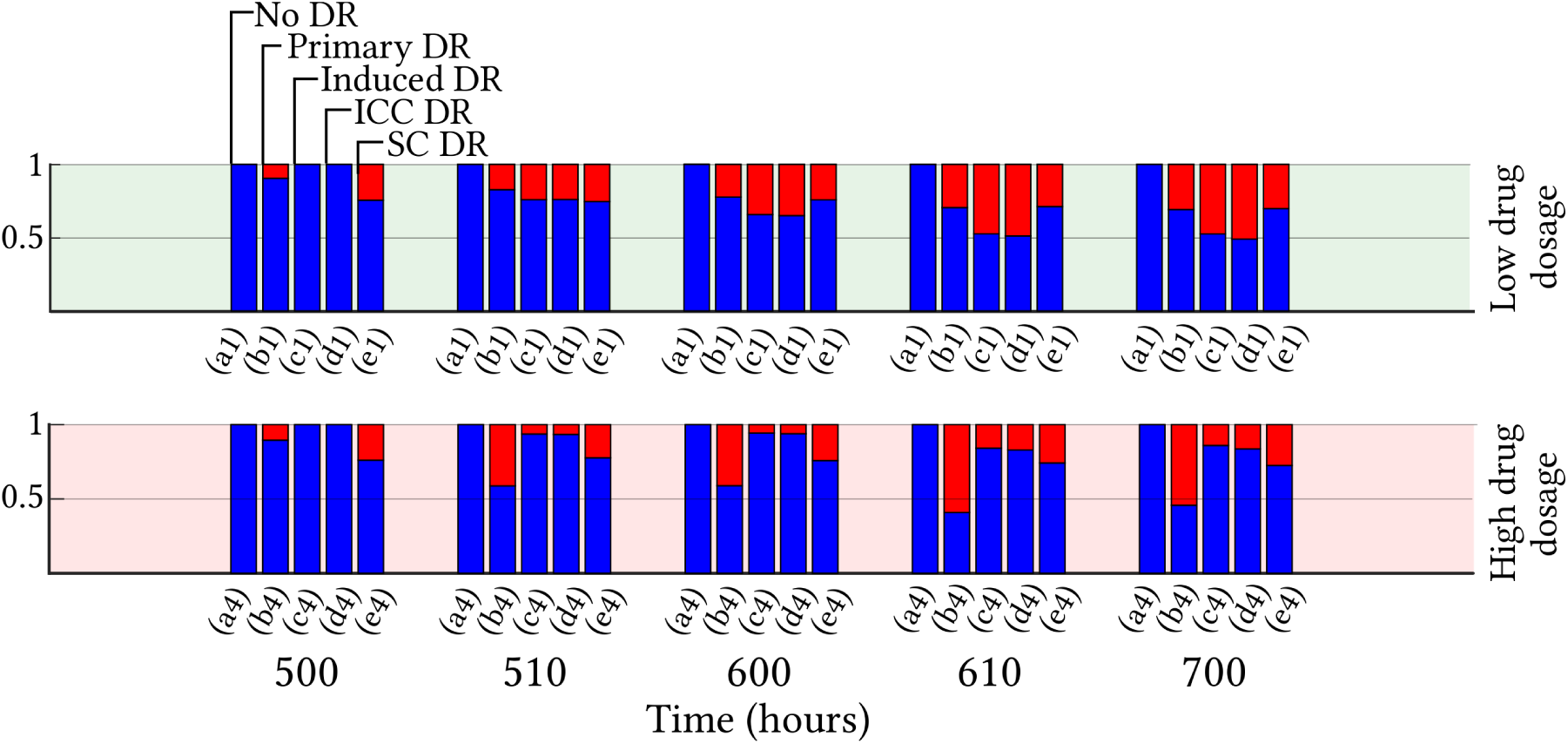
Diagram of the ratio between the drug sensitive subpopulation (blue) and the drug resistant sub-population. Various types of drug resistance are incorporated namely (a) No DR, (b) Primary DR, (c) Induced DR, (d) ICC DR, (e) SC DR. Results are shown at times 500 h (immediately before the first drug dose), 510 h, 600 h (immediately before the second drug dose), 610 h and 700 h (end of simulation) for low drug dosages, (1) 1C, and high drug dosage, (4) 8C according to the labelling in Table 2. Each diagram shows the average value based on 100 simulations.

The aim of this study is to qualitatively model drug response in cancer cell populations hosting drug resistant individuals. Drug resistance is here modelled from a collective of biological experiments, biological theory and clinical observations, and thus does not confer strictly with one cell line or one experiment. However, the developed *in silico* framework can be parametrised and calibrated appropriately for a cell-line specific study (as shown in a recent paper [8]), should relevant biological data become available in detail. Our *in silico* framework is equipped to handle various mechanisms concerning drug resistance, these mechanisms can be appropriately included or excluded in order to study a certain cell-line or a certain tumour scenario.

## 4 Conclusion

Enhanced chemotherapeutic drug resistance post-chemotherapy is an established clinical problem, this study provides insight into drug resistance and drug response in cancer cell populations on a cellular resolution. Our results show that, whilst chemotherapy is an effective way to reduce tumours, suboptimal drug dosages may contribute towards drug resistance and, by extension, tumour reinforcement. Thus, in accordance with Nietzschean philosophy, chemotherapy that does not kill a tumour may indeed make it stronger. Generally we found that drug resistance presenting independently of the drug, which thus may precede chemotherapy, is amplified by high drug dosages. However, drug resistance that is induced by drug presence is accelerated by low to moderate drug dosages. These findings are pictorially demonstrated in Figure 10.

**Figure 10:**
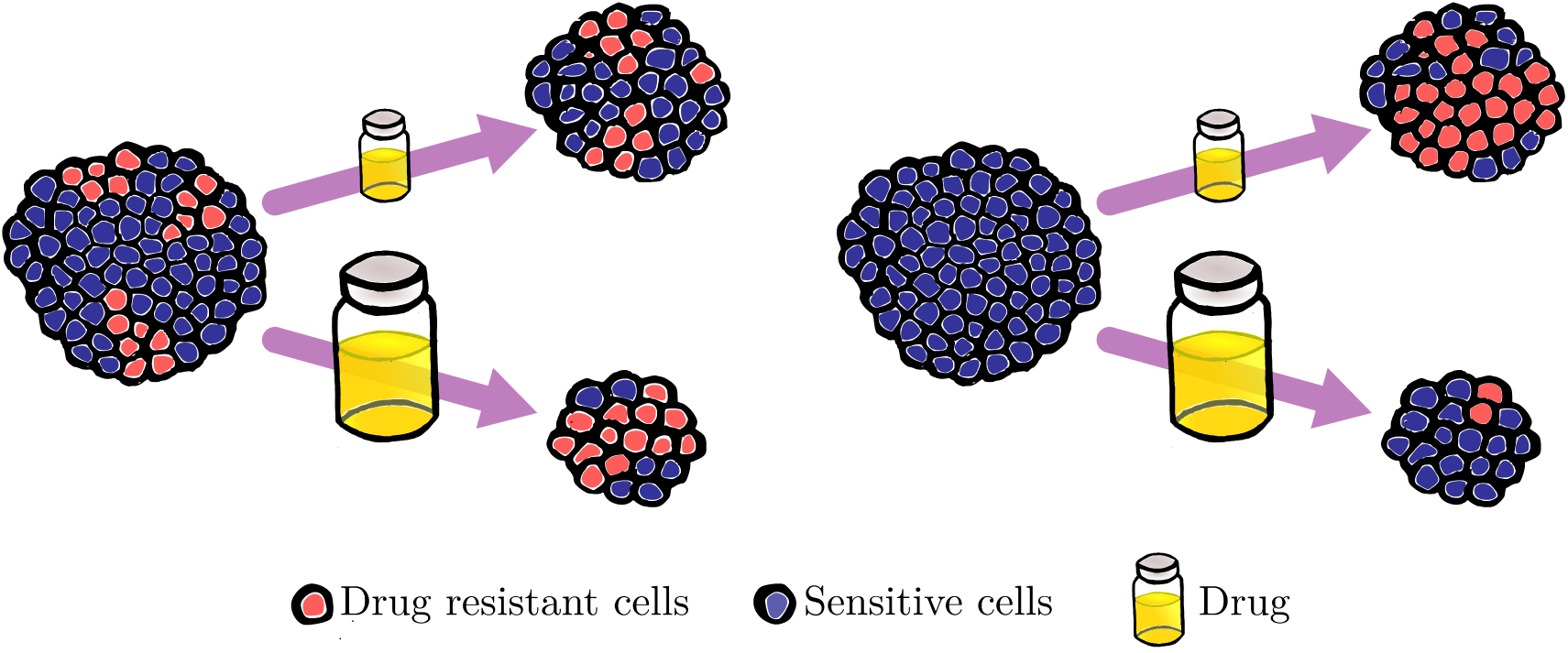
Tumours exposed to chemotherapeutic drugs given in low (top) or high (bottom) dosages, the latter dosage option kills more cancer cells. One of the tumours (left) expresses drug resistance pre chemotherapy, the other one (right) does not. Generally in our *in silico* experiment, drug resistance that occurs independently of the drug, and thus may precede chemotherapy, is amplified by high drug dosages (left). Conversely, drug resistance that is induced by drug presence is accelerated by low to moderate drug dosages (right).

## Acknowledgments

SH gratefully acknowledges the support of Swansea University PhD Research Studentship.

## Supplementary Material

### 1 Intracellular Dynamics

#### 1.1 Parameters involved in regulating the cell cycle

This section contains parameters occurring in Section 2.1.1, Equation 1 in the article. Parameters listed in Table 1 are gathered from a previous study by Powathil et al. [54].

**Table S1:** Parameters for the nondimensionalised form of Equation 1.

| Component | Rate constants ($\text{h}^{-1}$) | Dimensionless parameters |
| --- | --- | --- |
| [CycB] | $k_1 = 0.12, k'_2 = 0.12$ | $[HIF1] = \begin{cases} 1 & \text{if } K \leq 0.1 \\ 0 & \text{otherwise} \end{cases}$ |
| | $k''_2 = 4.5, [\text{p27/p21}] = 1.05$ | |
| [Cdh1] | $k'_3 = 3, k''_3 = 30, k_4 = 105$ | $J_3 = 0.04, J_4 = 0.04$ |
| [p55cdc<sub>T</sub>] | $k'_5 = 0.015, k''_5 = 0.6, k_6 = 0.3$ | $J_5 = 0.3, n = 4$ |
| [p55cdc<sub>A</sub>] | $k_7 = 3, k_8 = 1.5$ | $J_7 = 0.001, J_8 = 0.001, [\text{Mad}] = 1$ |
| [Plk1] | $k_9 = 0.3, k_{10} = 0.06$ | |
| [mass] | $\mu = \mu^+ + \epsilon \hat{\mu}$ | $\epsilon$ is randomised, $\epsilon \in [-1, 1]$ |

The value *m_*_* occurring in Equation (1f) denotes the maximum mass that a cell may reach, should it not divide. Following previous work by Tyson and Novak [73], here *m_*_* ≫ *max*([*mass*](*x,t*)), specifically *m_*_* = 10 · [*mass*](*x_nb_, t_nb_*), where [*mass*](*x_nb_*, *t_nb_*) corresponds to the mass of a “newborn” cell. The growth rate constants *μ*^+^ and 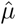 are here chosen to produce cell cycle lengths averaging 25 hours for drug sensitive cells that are well oxygenated, i.e cells that have the [*HIF*] component occurring in Equation 1a switched off (set to zero). For well oxygenated slow-cycling cells belonging to the implicitly drug resistant subpopulation SC DR, the average cell cycle length is roughly doubled to 50 hours. This is achieved by setting the growth rate components to be *μ*^+^ = *s ·* 0.01 and 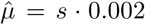, where *s* = 1 for sensitive, fast-cycling cells and *s* = 0.5 for slow-cycling, (SC DR) cells. Independently of drug resistant traits, the [*HIF*] component is activated (set to one) in hypoxic cells, this activation delays cell cycle progression and consequently yields a cell cycle length increase of approximately 20%.

#### 1.2 Details of the mathematical cell cycle model

Tyson and Novak have produced a series of papers which describe the underlying mechanisms of cell cycle progression and regulation in mathematical, implementable terms [43, 44, 73]. By recognising key proteins involved in controlling the cell cycle, they have managed to condense this complex biological phenomenon into a six-component regulatory molecular network [73]. This network can be translated into a system of ordinary differential equations, here Equations (1a) through to (1f) are gathered from one of their papers [73]. Appropriate proteins for mammalian cells are here used and changes in protein concentrations over time are computed in order to track cell cycle progression, where protein concentrations are measured in grams of protein per gram of total cell mass. In accordance with previous work by Powathil et al. [54], parameter values scaled to fit cell cycle lengths averaging 25 hours are used in order to match those of typical fast-cycling mammalian cells.

The opposing and oscillating nature between Cyclin-dependent protein kinases (Cdks) and the anaphase-promoting complex (APC) plays a central role in cell cycle regulation and yields a hysteris feedback loop. Cyclin B ([*CycB*]) is part of the Cdk family whilst *Cdh*1 is part of the APC. As is demonstrated in Figure S1a, the cell cycle control system consists of two steady states, namely the G1-phase state and the Syn-G2-M-phase state. [*CycB*] is low whilst [*Cdh*1] is high in the G1 state and conversely [*CycB*] is high whilst [*Cdh*1] is low in the Syn-G2-M state. Other auxiliary molecules are included in the model to enable appropriately lagging transitions between these two steady states, their concentrations over time are illustrated in Figure S1b. At the start of the cell cycle, the system will tend towards a G1 steady state and at the end of the cell cycle the system will tend towards a Syn-G2-M steady state. Details are provided by Tyson and Novak [73]. Cell mass, [*mass*], will double over the course of a cell cycle and is later reset, i.e halved, at cell division, as shown in Figure S1c. Now follows a description of each equation occurring in the system of ordinary differential equations used to regulate and track cell cycle progression, namely Equation (1a) through to Equation (1f).

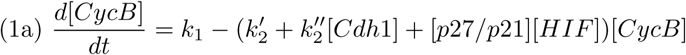

Cdks are necessary to replicate DNA, accordingly Cdk activity is high in the Syn-G2-M whilst Cdk activity is low in the G1 phase. The rate of change of [*CycB*] is governed by synthesis and reduction, where the reduction is both naturally occurring and induced by [*Cdh*1] presence. *p*21 and *p*27 are proteins that inhibit Cdks, here activated in hypoxic cells only via the [*HIF*] component, where *p*21 and *p*27 here merely are treated as parameters via the [*p*27/*p*21] factor [54].

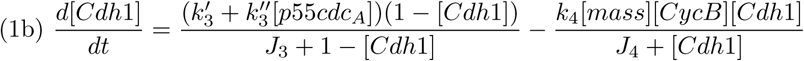

The p55cdc-APC in its total (*p*55*ctc_T_*) and active form (*p*55*cdc_A_*) is also part of the APC. The [Cdh1] time derivative obeys Michaelis-Menten type equations, where the occurring J-constants are Michaleis constants. Here [*CycB*] inhibits [*Cdh*1] activity whilst [*p*55*cdc_A_*] enhances it. The [*CycB*]-derived inactivation of [*Cdh*1] is assumed to occur in the cell nucleus, hence [*CycB*] will accumulate in the nucleus and it follows that [*CycB*] will increase with cell mass, thus the [*mass*] factor is incorporated in describing the suppressing effect that [*CycB*] has on [*Cdh*1]. For newborn cells, [*mass*] is low, however as the cell cycle progresses, [*mass*] will increases, consequently promoting [*CycB*] whilst demoting [*Cdh*1].

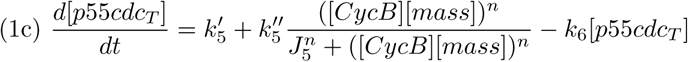

[*p*55*cdc_T_*] synthesises occurs naturally throughout the cell cycle but is also synthesised in the Syn-G2-M phase by [*CycB*], this is appropriately described by a Hill function. However p55cdc is not active once newly synthesised, this activation is described in Equation (1d).

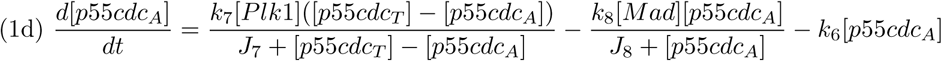

In the model, [*Plk*1] is included to transform p55cdc into its active form [*p*55*cdc_T_*]. Tyson and Novak [73] describe [*Plk*1] as a hypothetical enzyme driving [*p*55*cdc_T_*] activation and yielding a hysterical effect in the [*CycB*] activity. This effect is incorporated with Michalis-Mentin equations. [*MAD*] represents a family of checkpoint genes, here treated as a parameter, which are able to deactivate [*p*55*cdc_A_*], should DNA synthesis or chromosomes alignment not be completed rapidly enough to allow correct cell cycle advancement.

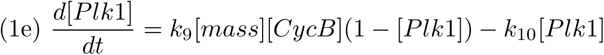

[*Plk*] decreases naturally whilst [*CycB*] enhances [*Plk*] activity, where again it is apparent that the [*mass*] component boosts [*CycB*] effects.

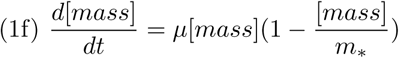

[*mass*] will double over the course of a cell cycle, following an adapted logistic equation. *μ* and *m*_*_ are described in 1.1.

**Figure S1:**
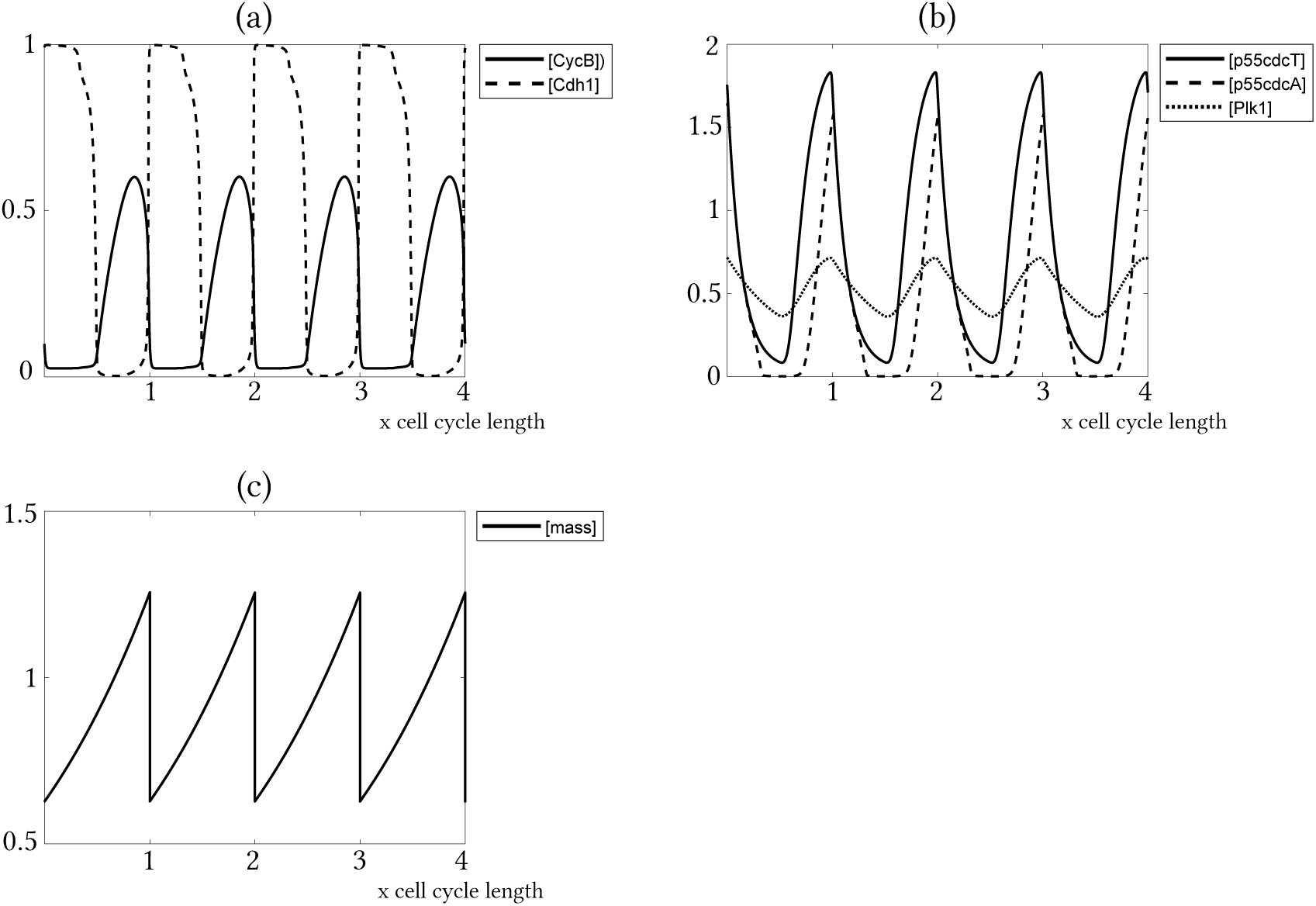
Time evolution of protein concentrations in one simulated cancer cell (measured in grams of protein per gram of total cell mass), where the cell cycle length is 25 hours. (a) The opposing dynamics between [*CycB*] and [*Cdh*1], which is key in the used mathematical cell cycle model. (b) Concentrations of the auxiliary proteins [*Pkl*1] and p55cdc in its total form [*p*55*cdc_T_*], and its active form [*p*55*cdc_A_*]. (c) Cell mass, [*mass*] over time, where [*mass*] is halved at the very start of each cell cycle, i.e at cell division.

### 2 Extracellular Dynamics

#### 2.1 Parameters for oxygen and drug distribution

This section contains parameters occurring in Section 2.1.2, specifically Equations 2 and 3.

**Table S2:** Parameters for Equations 2 and 3.

| Parameter | Symbol | Value | Reference |
| --- | --- | --- | --- |
| Oxygen diffusion coefficient | $D_k$ | $2.5 \cdot 10^{-5} [\text{cm}^2 \text{s}^{-1}]$ | [54] |
| Oxygen supply rate | $r_k$ | $8.2 \cdot 10^{-3} [\text{s}^{-1}]$ | [54] |
| Oxygen consumption rate | $\phi_k$ | $2 \cdot 10^{-1} [\text{s}^{-1}]$ | [54] |
| Cisplatin diffusion coefficient | $D_c$ | $7.6 \cdot 10^{-6} [\text{cm}^2 \text{s}^{-1}]$ | [54] |
| Cisplatin consumption rate | $\phi_c$ | 0 (*negligible*) | [54] |
| Cisplatin decay rate | $\eta_c$ | 1.316 $[\text{h}^{-1}]$ | *estimated* from [57] |

Diffusion and production rates of oxygen and drugs vary across the grid so that [54]

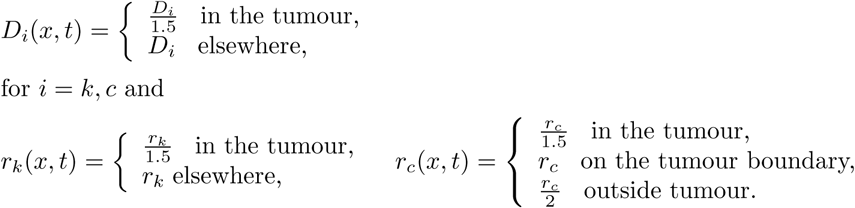

The drug decay rate *η_c_* is estimated to match the half-life time of cisplatin which has been reported as 31.6 ± 6 minutes [57]. Cisplatin supply is here modelled as instantaneous and thus equal to *r_c_* at the two time steps conferring with drug administration (t= 500 hours, t= 600 hours) and zero for all other time steps. *r_c_* is estimated and nondimensionalised, in absolute form the total drug dosage 1C takes roughly the same value as the absolute oxygen concentration (see Section 2.2 and Table S3) at drug administration time. Thus we assume oxygen and drug levels to be comparable at the time of drug administration. The cisplatin supply rate *r_c_* is scaled according to the chosen drug dosage so that

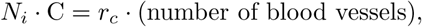

hence *r_c_* corresponds to the amount of drug produced at one blood vessel cross section at one time step. Here *i* = 0,1, 2, 3,4 and *N_i_* = 0,1, 2,4, 8 in accordance with the possible drug dosages explored in the *in silico* experiment which are 1C, 2C, 4C and 8C. A cell in location *x* at time *t* is killed if the drug concentration is such that 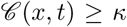. C and *κ* are parametrised so that 1C kills half of the cell population in absence of any included drug resistant mechanisms. Specifically here C= 10^4^ (equal to the number of grid points) and *κ* = 0.18. Since C is equal to the number of grid points, all cancer cells would die immediately from drug exposure if drugs were produced homogeneously across the grid, as such a scenario would yield a drug concentration 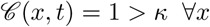.

#### 2.2 Spatial oxygen distribution

The total, absolute amount of oxygen in the system fluctuates over time due to the chosen oxygen equation, parameters and boundary conditions. However scaled oxygen values are used in order to evaluate spatial oxygen distribution and determine hypoxia, these values are re-scaled at every time step [52]. In Figure S2 cell-maps and oxygen-maps demonstrating spatial oxygen distribution at certain times are provided, these maps are visualised using ParaView [5]. In Table S3, various oxygen measurements for certain time points are listed in both absolute and scaled forms. Here 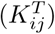 denotes the absolute oxygen value in grid point (*i, j*) at time *T*, and similarly 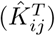 denotes the scaled oxygen value in grid point (*i, j*) at time *T*. Here *i* and *j* are spatial integer indices ranging from 1 to 100 as a square grid with 100^2^ grid points is used. For absolute oxygen values these listed measurements are specifically; the total amount of oxygen in the system 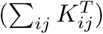, the average oxygen value at one grid point 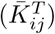, the oxygen value at the grid point with the maximum amount of oxygen 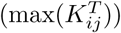 and the oxygen value at the grid point with the minimum amount of oxygen 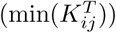. The same measurements are done for scaled oxygen values, using the hat notation to denote that the values are scaled.

**Figure S2:**
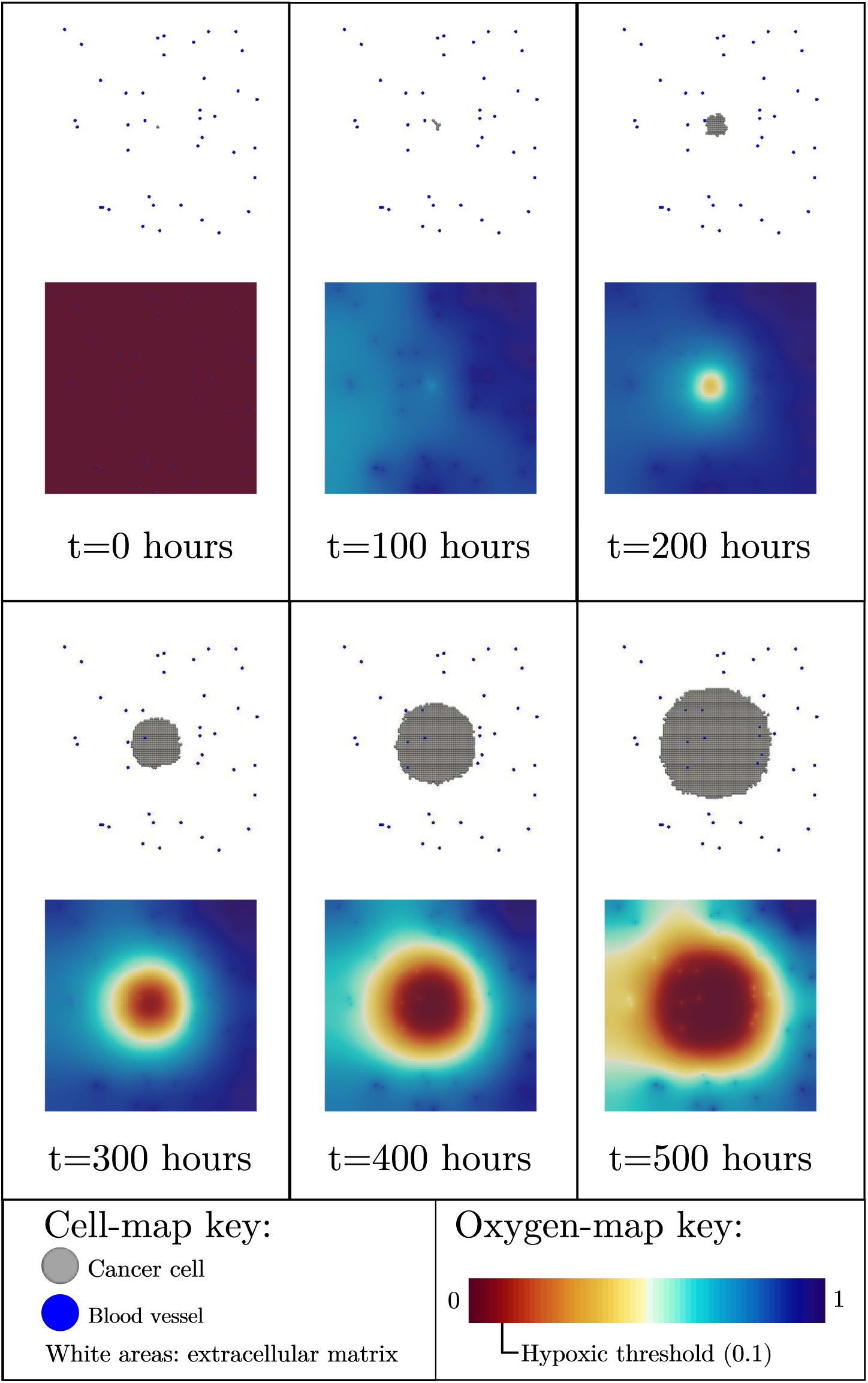
Cell-maps (top) and oxygen-maps (bottom) at certain times pre chemotherapy administration. The oxygen-maps demonstrate the spatial oxygen distribution in terms of scaled, nondimensionalised oxygen values ranging between 0 and 1 at each grid point.

**Table S3:** Nondimensionalised oxygen values of the system at certain time points, in both absolute form and scaled form (hat notation).

| time [h] | $\sum_{ij} K_{ij}^T$ | $\bar{K}_{ij}^T$ | $\max(K_{ij}^T)$ | $\min(K_{ij}^T)$ | $\sum_{ij} \hat{K}_{ij}^T$ | $\bar{\hat{K}}_{ij}^T$ | $\max(\hat{K}_{ij}^T)$ | $\min(\hat{K}_{ij}^T)$ |
| --- | --- | --- | --- | --- | --- | --- | --- | --- |
| 0 | 0.41 | 0 | 0.01 | 0.00 | 40.05 | 0.00 | 1.00 | 0.00 |
| 100 | 41048.10 | 4.10 | 4.92 | 3.50 | 8334.76 | 0.83 | 1.00 | 0.71 |
| 200 | 70535.20 | 7.05 | 8.30 | 3.36 | 8498.16 | 0.85 | 1.00 | 0.40 |
| 300 | 58273.21 | 5.83 | 7.75 | 0.69 | 7514.37 | 0.75 | 1.00 | 0.09 |
| 400 | 29163.37 | 2.92 | 4.74 | 0.09 | 6155.72 | 0.62 | 1.00 | 0.02 |
| 500 | 12093.96 | 1.21 | 2.62 | 0.02 | 4619.06 | 0.46 | 1.00 | 0.01 |

### 3 Drug Resistance

This section contains parameters and algorithms used in Sections 2.2.1 through to 2.2.4 in the original paper. The chosen parameter values corresponding to drug resistance for the *in silico* framework are listed in Table S4, where in hypoxic regions *α_ex_* is increased to *2α_ex_*. The results in this study are qualitative, when varying the parameters in Table 3, as done in Supplementary Material Section 4.2, the obtained qualitative results are robust in regards to chemotherapy response. *τ* is chosen to be 30 minutes as this is close to the half-life time of cisplatin. *α_pri_, χ_ind_, α_ex_* and *α_SC_* are parametrised to be low and yield approximately the same ratio between the sensitive subpopulation and the drug resistant subpopulation across the different tests (b1), (c1), (d1) and (e1) 10 hours after the first drug administration, as demonstrated in Figure 9 in the article. This is in order to allow for easy comparisons, in regards to drug response evaluations, amongst tests.

**Table S4:** Parameters concerning drug resistance used in the *in silico* framework.

| $\alpha_{pri}$ | $\chi_{ind}$ | $\tau$ | $\alpha_{ex}$ | $\alpha_{SC}$ |
| --- | --- | --- | --- | --- |
| 0.01 | $\kappa/10$ | 30 minutes | 0.07 | 0.07 |

#### 3.1 Model

This section demonstrates partial algorithms which are implemented to compute if a cell is drug resistant or not.

##### 3.1.1 Primary drug resistance

Each time cell division occurs there is a chance that a drug resistant cell will be produced. In the model, when cell division occurs it is said that a mother cell produces a daughter cell. If the mother cell is drug resistant (DR), the daughter cell will inherently be drug resistant. However, if the mother cell is sensitive (S), the daughter cell may or may not be drug resistant according to a stochastic model. At each cell division a value *α* ∈ [0,1] is randomised, if *α* ≤ *α_pri_* then the daughter cell is drug resistant, otherwise it is sensitive, as schematically shown in Figure 3.1.1.

**Figure S3:**
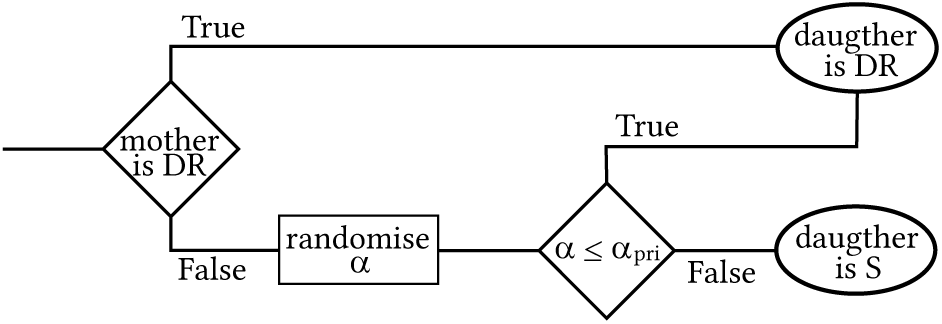
Partial algorithm for determining primary drug resistance.

##### 3.1.2 Induced drug resistance

If a cell has experienced a minimum drug concentration *χ_ind_* for *τ* time units, drug resistance is induced in the cell. Thus in the model each cell has its own counter, counter*_n_*, which increments each time step that cell*_n_* experiences a drug concentration 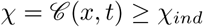. This is schematically illustrated in Figure 3.1.2. If drug resistance has been produced in a mother cell it produces drug resistant daughter cells.

**Figure S3:**
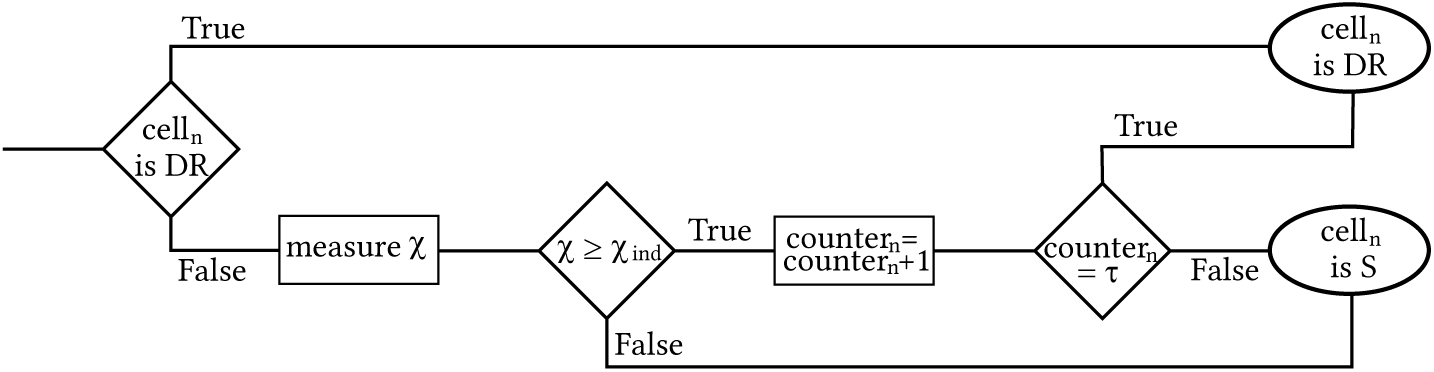
Partial algorithm for determining induced drug resistance.

##### 3.1.3 Communicated drug resistance via exosomes

Once per cell cycle, each cell that has acquired drug resistance by induces drug resistance (see the above section) or communicated drug resistance (as described in this section) has a chance *α_ex_* of producing and secreting an exosome. In the model, a value *α* ∈ [0,1] is randomised, if *α* ≤ 2*α_ex_* in hypoxic regions, or *α* ≤ *α_ex_* in normoxic or hyperoxic regions, an exosome is produced and sent off in a random direction. The first sensitive cell that the exosome hits becomes drug resistant, as schematically shown in Figure 3.1.3. Mother cells that have acquired drug resistance via exosomes pass on their drug resistance to daughter cells.

**Figure S3:**
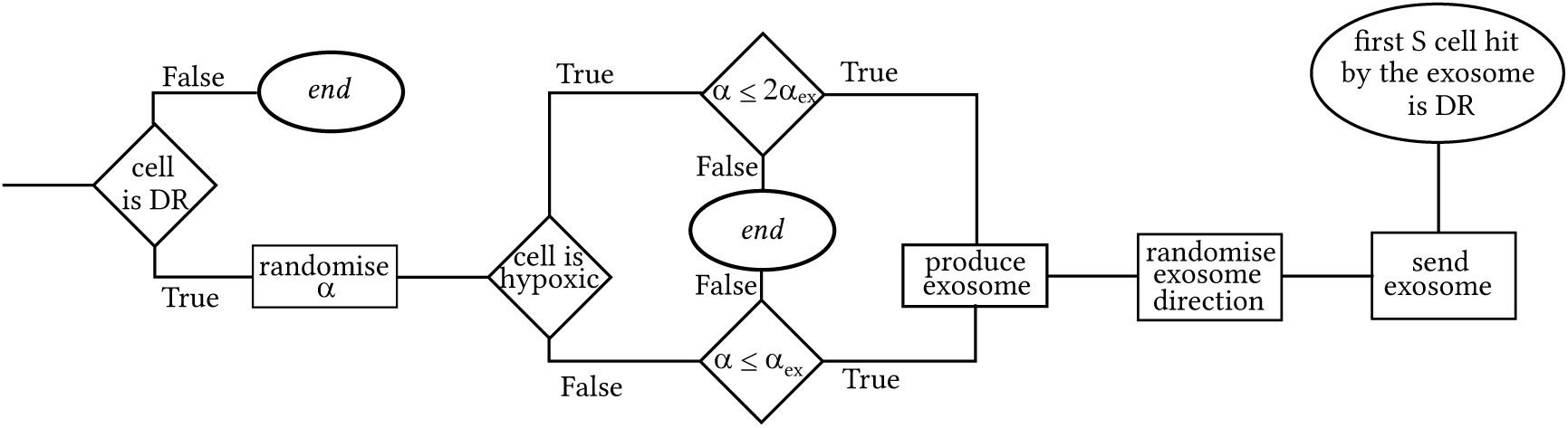
Partial algorithm for determining exosome production.

##### 3.1.4 Cell-cycle mediated drug resistance by slow-cycling cells

Once per cell cycle there is a chance that a sensitive, fast-cycling (FC) cell will spontaneously convert into a slow-cycling (SC), drug resistant state. To check if such conversion occurs, *α* value *α* ∈ [0,1] is randomised. If *α* ≤ *α_SC_* then the cell converts, otherwise it does not, as demonstrated in Figure 3.1.4. Slow-cycling (SC DR) mother cells yield slow-cycling (SC DR) daughter cells.

**Figure S3:**
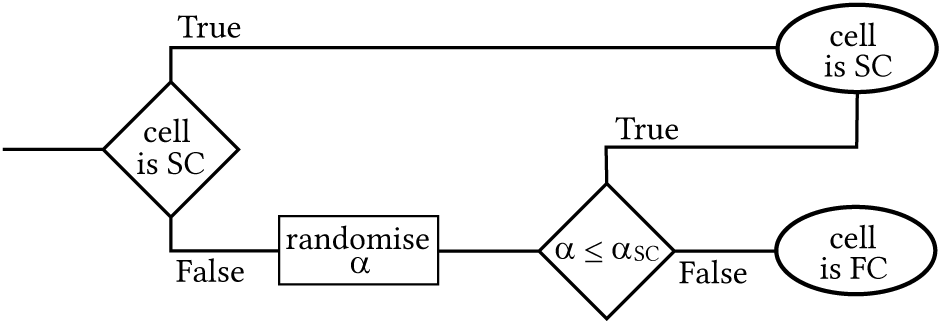
Partial algorithm for determining the spontaneous convertion from a slow-cycling to a fast-cycling state.

### 4 Sensitivity Analysis

In this section, sensitivity analysis is performed on the main results presented in the article, specifically on the results demonstrated in Figure 4. In Section 4.1, statistical data concerning these tests are provided numerically and in Section 4.2 robustness tests are performed in order to confirm that the qualitative results obtained in this study are robust to variations of chosen and estimated parameters.

#### 4.1 Statistical data

This section provides listings of the mean value (mean), standard deviation (S.D) and 95 %-confidence interval (I.C(95%)) for test results (a1) through to (e4), presented by graphs in Figure 4. Here all values have been rounded to integers, corresponding to full number of cells.

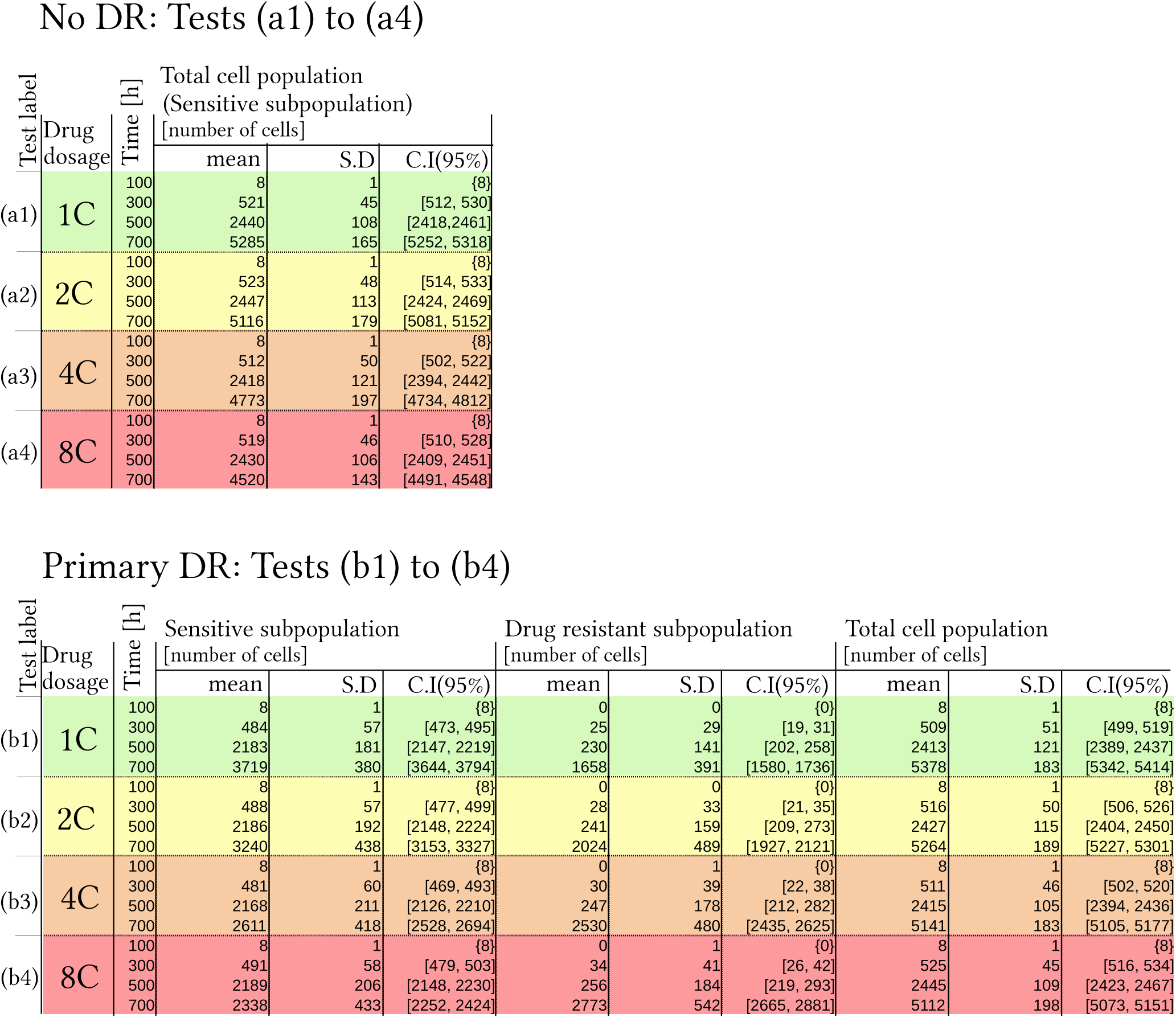
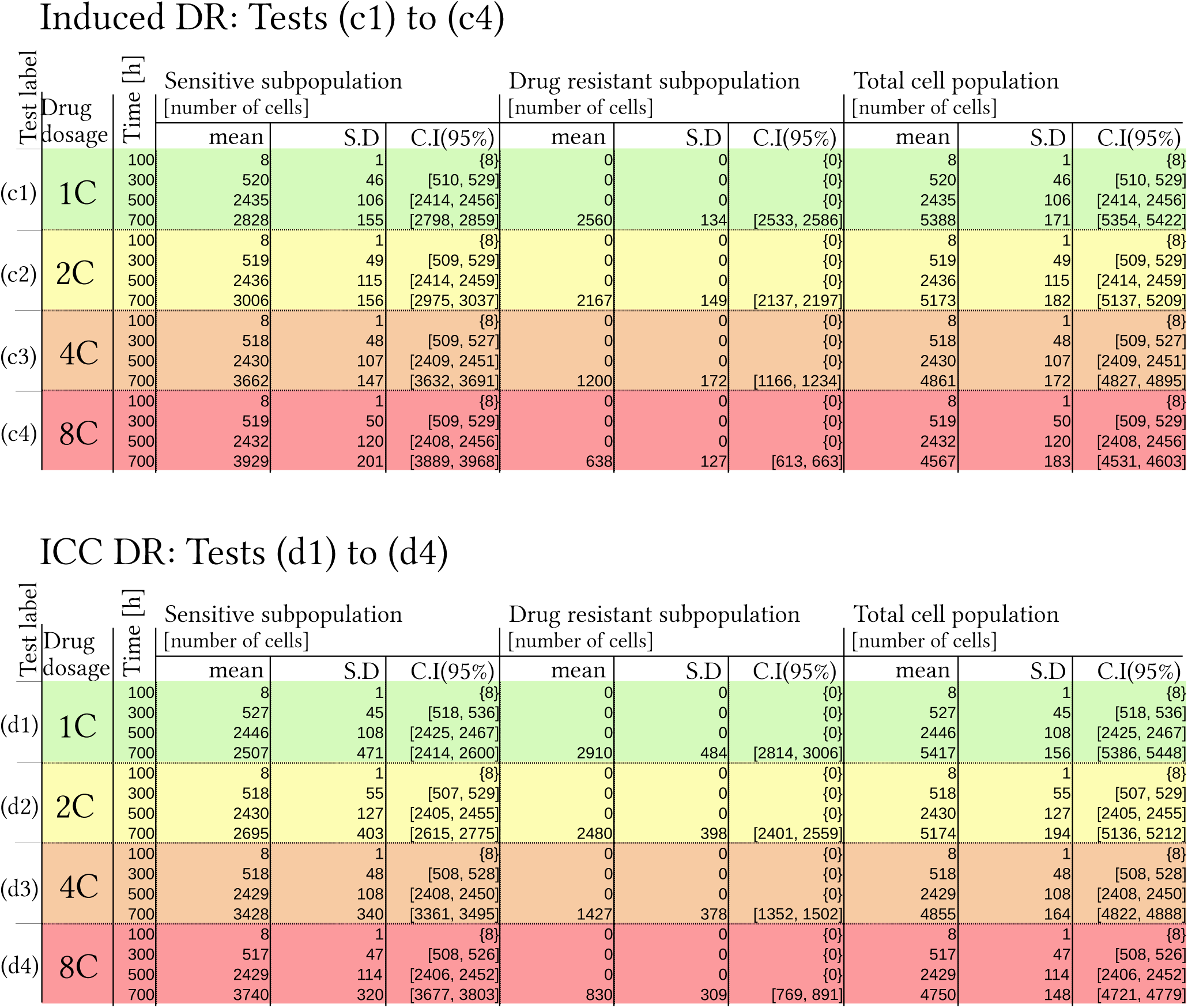
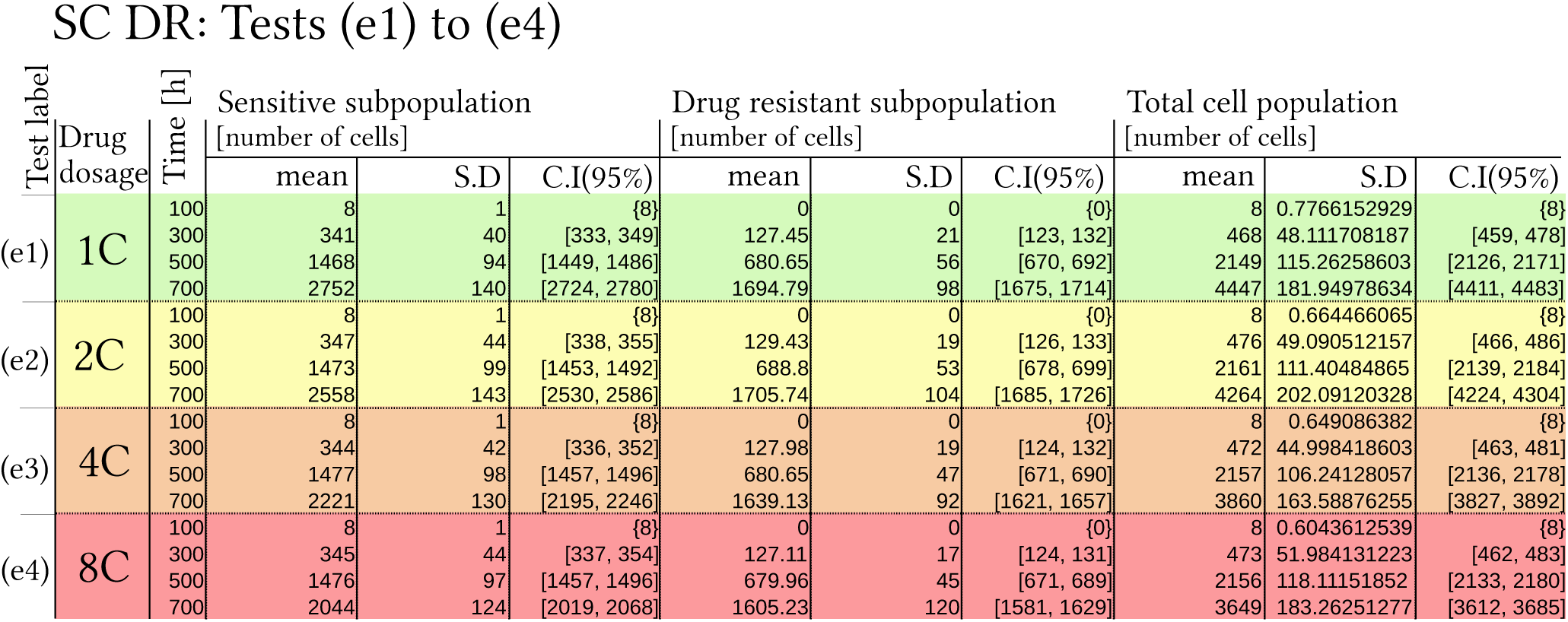

#### 4.2 Parameter variation

To verify that our results are robust in regards to the chosen parameters listed in Table S4, a sensitivity analysis is preformed in which parameters are varied, one at a time, according to Table S5. Each such sensitivity test is performed 100 times, results are provided in Figure S7. These results show that our qualitative findings, concerning drug response in cancer cell populations hosting various types of drug resistance, hold for parameter variations. Indeed the ratio of drug resistant cells increases with high drug dosages in cases where drug resistance precedes chemotherapy, here in experiments (b) Primary DR and (e) SC DR. Conversely drug-induced drug resistant subpopulations are promoted in scenarios with low drug dosages, here in experiments (c) Induced DR and (d) ICC DR.

**Table S5:**
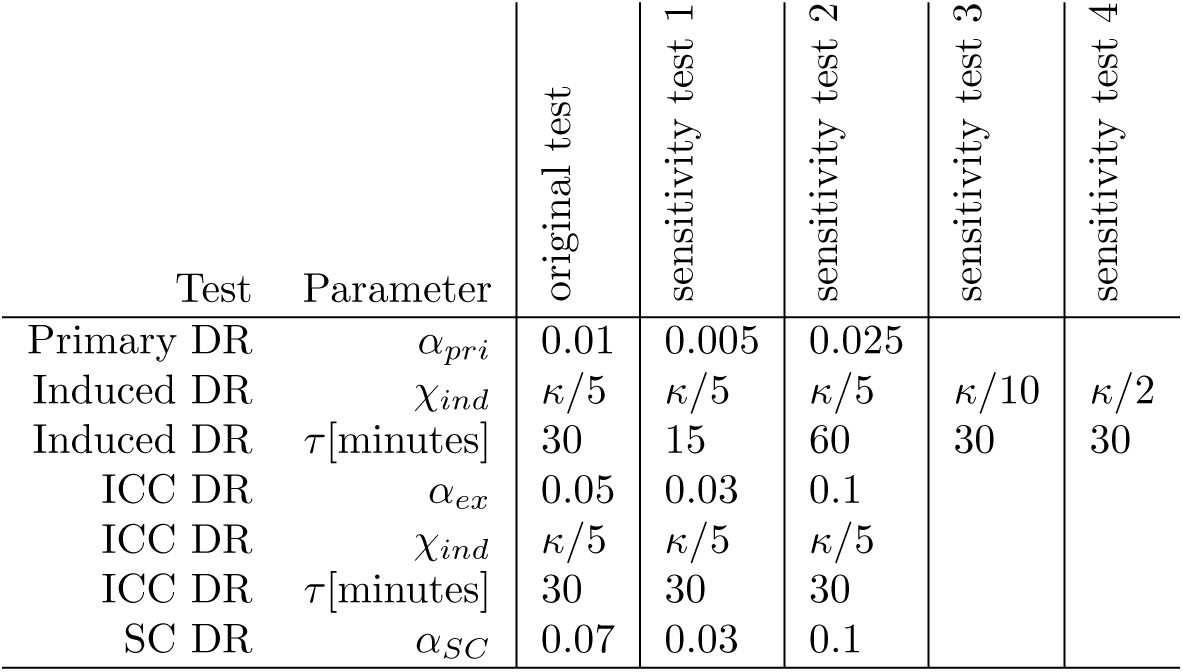
Parameters used in testing the robustness of the model.

| Test | Parameter | original test | sensitivity test 1 | sensitivity test 2 | sensitivity test 3 | sensitivity test 4 |
| --- | --- | --- | --- | --- | --- | --- |
| Primary DR | $\alpha_{pri}$ | 0.01 | 0.005 | 0.025 | | |
| Induced DR | $\chi_{ind}$ | $\kappa/5$ | $\kappa/5$ | $\kappa/5$ | $\kappa/10$ | $\kappa/2$ |
| Induced DR | $\tau$[minutes] | 30 | 15 | 60 | 30 | 30 |
| ICC DR | $\alpha_{ex}$ | 0.05 | 0.03 | 0.1 | | |
| ICC DR | $\chi_{ind}$ | $\kappa/5$ | $\kappa/5$ | $\kappa/5$ | | |
| ICC DR | $\tau$[minutes] | 30 | 30 | 30 | | |
| SC DR | $\alpha_{SC}$ | 0.07 | 0.03 | 0.1 | | |

**Figure S4:**
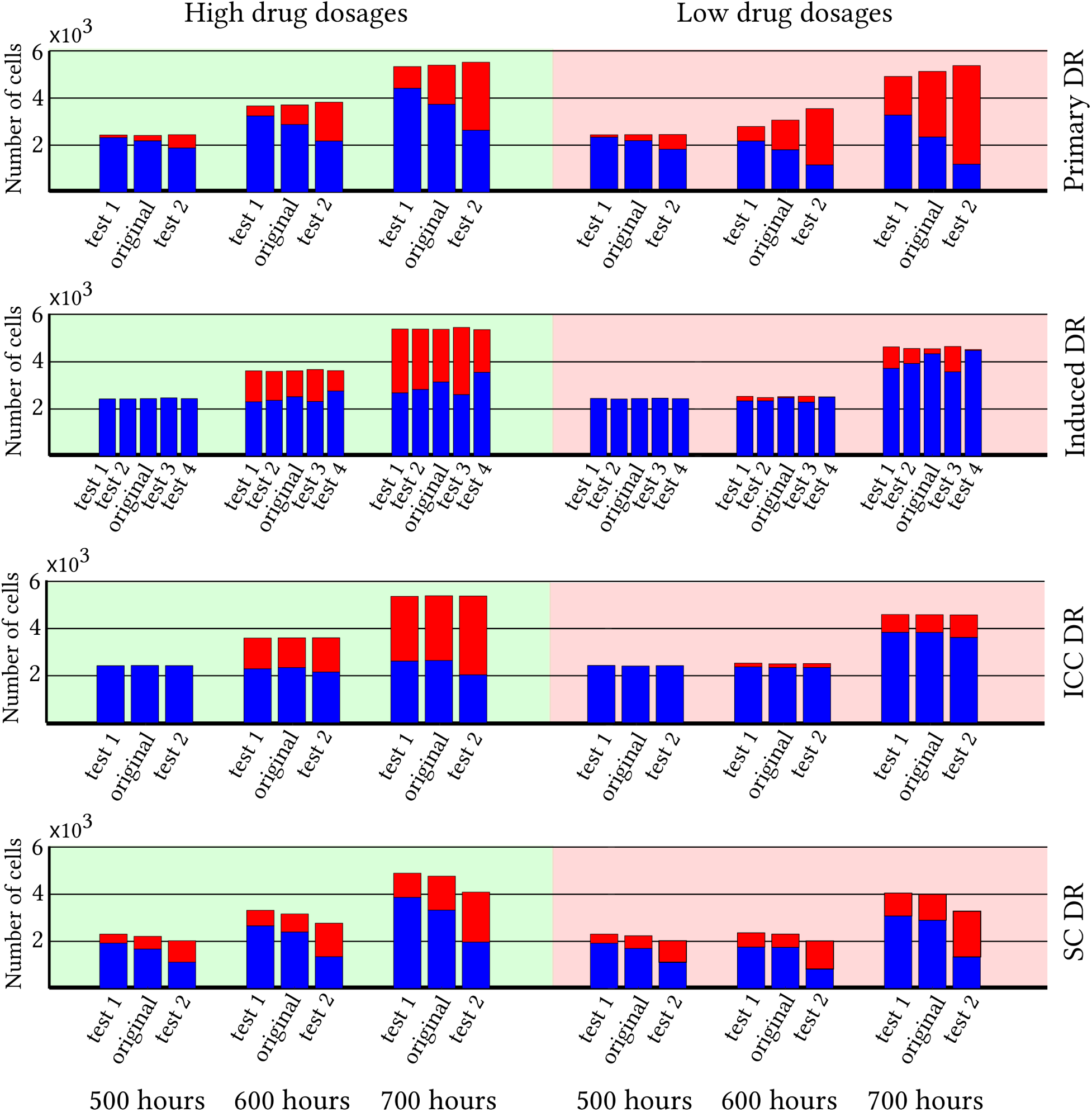
Sensitivity analysis, showing the number of sensitive (blue) and drug resistant (red) cells at three time points when low (left) and high (right) drug dosages are administered, namely 1C and 8C respectively. Each test is performed 100 times and the parameters used in each test are listed in Table S5. Thus for Primary DR, only the parameter *α_pri_* is varied. For Induced DR, *χ_ind_* and *τ* are both varied, one at a time, according to Table S5. For ICC DR only *α_ex_* is varied and similarly for SC DR only *α_SC_* is varied.

